# Improved cryo-EM reconstruction of sub-50 kDa complexes using 2D template matching

**DOI:** 10.1101/2025.09.11.675606

**Authors:** Kexin Zhang, Timothy Grant, Nikolaus Grigorieff

## Abstract

Visualizing the structures of small proteins and complexes has been a longstanding challenge in single-particle cryo-EM. Some of these targets have been successfully resolved by binding to antibody fragments (Fabs) or fusing with external scaffolds to increase their size. Recent advances in conventional single-particle techniques have enabled the determination of an increasing number of structures smaller than 100 kDa, achieving resolutions relevant to drug research. Compared to X-ray crystallography, cryo-EM preserves the near-native states of biomolecules, can resolve structural heterogeneity, and has the potential to apply to a wide range of targets. In this work, we demonstrate that the alignment and reconstruction of small macromolecular complexes can be improved using high-resolution structures as priors combined with 2D template matching. Using this method, we reconstructed a previously intractable ∼ 43 kDa protein kinase and improved the density of its ligand-binding site. Our theoretical analysis predicts that this method can further extend single-particle cryo-EM to important drug-binding complexes well below 50 kDa. Cryo-EM | 3D reconstruction | signal detection | template matching _61_

## Introduction

Since the “resolution revolution” in 2014 (1), single-particle cryo-electron microscopy (cryo-EM) has become widely used, emerging as a powerful alternative, and in some cases surpassing X-ray crystallography for structurally challenging samples. In particular, the 1.2 Å reconstruction of apoferritin, where individual hydrogen atoms are resolved, demonstrates that single-particle cryo-EM has now reached true atomic resolution for well-defined systems (2, 3). Despite recent advances in single-particle cryo-EM, structure determination for sub-50 kDa complexes remains challenging. Among EM structures deposited in the Protein Data Bank (PDB) since 2015 with resolutions better than 4 Å, 97% correspond to macromolecules larger than 50 kDa (4). Small complexes contain fewer atoms and scatter fewer electrons, leading to images with lower signal-to-noise ratios (SNRs). Particle alignment, typically based on calculating the crosscorrelation coefficients between the particle and reference (5), depends on the amount of phase contrast (signal) in the image. The weak contrast of small complexes against background noise makes it difficult to align particles accurately to calculate a high-resolution reconstruction. Moreover, single-particle experiments typically apply a total exposure of 30-50 electrons per Å^2^, leading to accumulating radiation damage. The diffusion of small particles due to beam-induced motion may further reduce the alignable high-resolution signal (6).

Extending single-particle cryo-EM to sub-50 kDa targets will open new avenues for studying critical drug-binding interactions and advance structure-based drug discovery. Approximately 75% of proteins in the human proteome are below 50 kDa (7). While X-ray crystallography is not inherently limited by small molecular weight, it requires well-ordered crystals, which are often difficult to obtain. Solution and solid-state NMR spectroscopy, on the other hand, provide atomic-level information derived from observables such as chemical shifts. However, these methods are generally limited to smaller particles, require highly concentrated samples, and data analysis is laborious, making them less suitable for high-throughput studies (8–10). Single-particle cryo-EM offers unique benefits compared to these two techniques, producing rich structural data in the form of images and enabling the visualization of biomolecules in near-native states at high resolution without the need for crystallization. To overcome the lower size barrier in cryo-EM, many strategies focus on increasing the apparent size and rigidity of small particles by binding them to external scaffolds or antibodies (11–14). However, finding good scaffolds or antibodies is difficult, and they sometimes create structural artifacts (15). Thus, there remains an incentive to develop approaches that enable high-resolution structure determination of isolated sub-50 kDa complexes in their near-native conformations.

Theoretical estimates of the lower molecular weight limit, dating back to 1995, suggested that a 3D reconstruction at 3 Å resolution is possible for particles as small as 38 kDa, given that ∼ 12,600 images are averaged (16). This calculation assumed perfect images, a perfect reference to align the particle images, and a total exposure limited to 5 electrons Å^−2^. Later, using the Rose criterion, a more optimistic prediction of 17 kDa was calculated, requiring only one-ninth as many images to be averaged (17). Since those early calculations, single-particle cryo-EM has seen significant technical advancements. The introduction of direct electron detectors (DEDs), combined with exposure weighting, now allows much higher electron exposures by recording data as movies and improving image resolution through motion correction (18, 19). More recently, the development of a laser phase plate offers the potential to further improve image contrast by using a high-intensity laser beam to introduce a stable and tunable phase shift to enhance low-resolution features in samples with weak contrast such as small complexes (20–23). Additionally, cooling specimens at liquid-helium temperatures can delay radiation damage and reduce information loss during imaging (24). Recent work has shown that using gold specimen support with 100 nm diameter holes at liquid-helium temperatures allowed imaging with better quality compared to liquid-nitrogen temperatures (25).

2D template matching (2DTM) was previously developed to identify particles in cellular cryo-EM images using a high-resolution template with high accuracy (26–28). In a 2DTM search, cross-correlation coefficients are calculated between the image and the template to determine the targets’ location and orientation. Previous work from our lab has shown that molecular features not in the template can be reconstructed from 2DTM-derived targets with high resolution (29). Building upon this, in this paper, we show that the alignment of sub-50 kDa complexes can be improved using 2DTM and stringent particle selection. We apply this method to a previously published dataset of the ∼ 43 kDa catalytic domain of protein kinase A (30) and demonstrate improved reconstruction of its ligand-binding sites. We develop a theoretical framework showing that the lower molecular weight limit can be further reduced to approximately 7.1 kDa with an exhaustive five-dimensional search, and to 5.7 kDa with a constrained search, assuming phase plate and liquid-helium cooling are used. Our findings highlight the potential of 2DTM to expand the applicability of cryo-EM to a broader range of biologically and pharmaceutically important targets, paving the way for structural studies of small complexes that remain difficult for standard workflows.

## Results

### Recovery of densities omitted from the template in a 43 kDa protein kinase

We evaluated the ability of 2DTM to recover omitted ligand densities by performing a template matching search and reconstruction with specific features omitted from the template. We used the published dataset of a 42.8 kDa protein kinase (EMPIAR-10252) (30). Particle stacks were generated from single-particle data using coordinates output from 2DTM searches with templates missing certain structural components. A subset of particles was then selected based on 2DTM statistics and image quality measurement from fits of the contrast transfer function (CTF) of the micrographs these particles came from. The selected particle stack was used for 3D reconstruction. This “omit-template” strategy follows the *baited reconstruction* approach (29) and was designed to detect template bias in reconstructions calculated from targets detected by 2DTM. Here, we use it to show that we can reconstruct the density of small ligands that are not included in the template but are bound to the imaged molecules in the cryo-EM dataset. We explored a range of template deletion scenarios: from omitting ATP and residues on an alpha-helix, to removing the binding pockets around ATP at various radii. Below, we detail how each deletion strategy affected the final density map. Consistently, we found that ATP density was robustly recovered even when ATP was deleted from the 2DTM search template. These results demonstrate that 2DTM can recover ligand densities, providing biological insights into ligand binding and flexibility. Because the corresponding features were omitted from the search template, the recovered densities cannot result from direct inclusion of those features in the template (29).

#### Deleting ATP, inhibitor, and an alpha-helical turn

We used the 2.2 Å X-ray model (PDBID: 1ATP, (31)) to generate a high-resolution 3D template in which the ATP, two Mn^2+^ ions, and six alpha-helix residues (residues 222–227) were deleted from the model. The final template, with a molecular weight of 38.0 kDa (non-hydrogen atoms only), was simulated using the `simulate` program in *cis*TEM with a uniform B-factor of 30 Å^2^ and a pixel size of 1.117 Å/pixel (32). Using the previously developed 2DTM p-value for particle picking and angular alignment (33), we performed 2DTM searches at Nyquist resolution and generated an initial stack of candidate particles, which were then subjected to further selection using 2DTM-derived statistics and CTF fitting parameters. A final stack of 7,353 particles was used for 3D reconstruction in *cis*TEM. Figure 1(a) compares the cryo-EM maps from the single-particle reconstruction (30), the 2DTM template, and the 2DTM-based reconstruction. The angular distribution of the 2DTM-derived particle stack and the Fourier Shell Correlation (FSC) curve are shown in Figure 1b. Because particles were aligned to a single high-resolution template during 2DTM, the reported 2.6 Å resolution is **not a gold-standard** FSC and does not reliably reflect true map quality. Figure 1c provides a close-up view of the ATP-binding site and the deleted residues, with the X-ray model overlaid for comparison. The average Q-score between the 2DTM reconstruction and the X-ray model was calculated using the `MapQ` command line tool (34, 35). Q-scores approaching 1 suggest atomic resolution, where individual atoms are resolved. Values near 0.5 indicate visible side chains while scores around 0.2 reflect unresolved side chains but resolved secondary structure (35). The Q-score for ATP was 0.60, indicating good agreement between the 2DTM reconstruction and the X-ray model, while the two Mn^2+^ ions had Q-scores of 0.48 and 0.73. The Q-scores of deleted residues 222–227 (Trp, Ala, Leu, Gly, Val, Leu) were 0.63, 0.51, 0.57, 0.54, 0.61, and 0.53, respectively. Additionally, the shape of the alpha-helix turn is clearly resolved, despite being absent in the template. The density of these features was recovered despite their absence from the search template. To further validate the recovered density, we performed Phenix real-space grouped occupancy refinement of the full 1ATP model against the omit reconstruction. Omitted residues 222–227 refined to occupancies of 0.55–0.80 (mean 0.72) and ATP to 0.61, while template-included control residues 150–155 remained near 1.0 (mean 0.96), supporting partial recovery of density in the omitted regions (Supplementary file 1).

**Figure 1.**
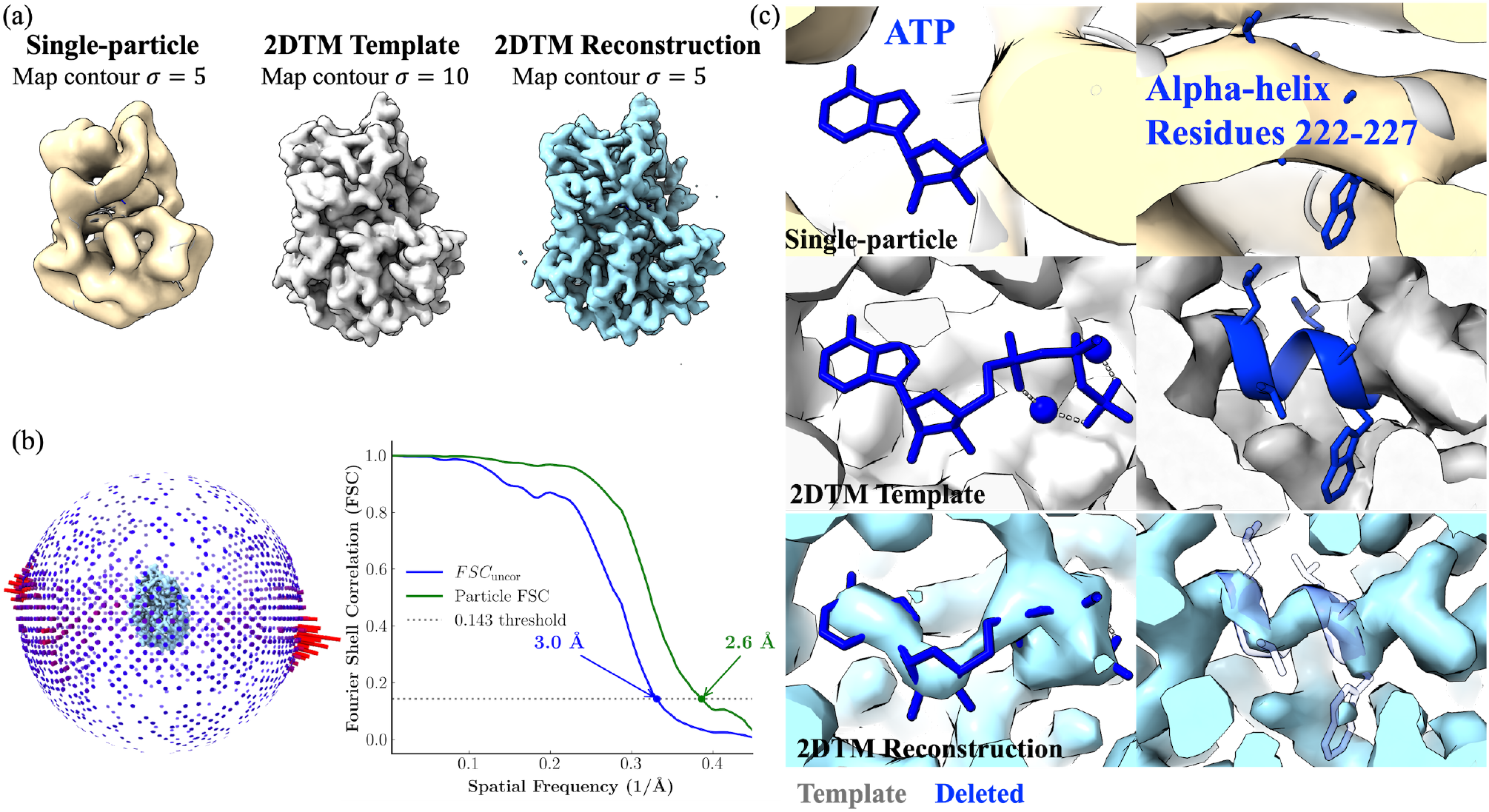
3D reconstruction of protein kinase using 2DTM-derived particle stack. **(a)** From left to right: the single-particle reconstruction (EMDB:0409) (30), the 2DTM template, and the 2DTM-derived reconstruction of the protein kinase. **(b)** Angular distribution plot and FSC curve calculated using *cis*TEM. Note that these FSC curves are not gold-standard FSCs, as the reconstruction uses orientations determined by 2D template matching rather than independent half-set refinement. *FSC*_uncor_ denotes the uncorrected FSC computed within a generous mask, which crosses the 0.143 threshold at 3.0 Å, and “Particle FSC” denotes the solvent-corrected FSC obtained using the mask-volume correction factor as described in the *cis*TEM/Frealign framework (64). **(c)** Densities at the ATP-binding site and deleted residues 222–227. The X-ray structural model (PDBID: 1ATP) (31) is shown with atoms retained in the template colored grey and deleted atoms colored blue. The alpha-helical model for residues 222–227 is rendered transparent to better show the recovery of the helical turn in the reconstruction. The average Q-scores between the 2DTM reconstruction and the X-ray model were calculated using the `MapQ` command line tool (34, 35). Q-scores for ATP, Mn^2+^, and deleted residues 222–227 were 0.60, 0.48/0.73, 0.63, 0.51, 0.57, 0.54, 0.61, and 0.53, respectively.

#### Robust reconstruction of the ATP binding pocket

To test the robustness of 3D reconstruction as more atoms are excluded from the template, we performed a series of 2DTM searches and reconstructions using templates with residues deleted within specified radii of the bound ATP. In Figure 2, we show the results of three different templates. We tested spherical deletion radii of 3.0 Å (Figure 2a and b) and 5.5 Å (Figure 2c) centered on the ATP position in the X-ray model. In addition to the ATP binding pocket, we also deleted Mn^2+^ in all templates and furthermore deleted IP20, an inhibitory pseudo-peptide substrate, in Figure 2b. The molecular weights of templates and numbers of particles in the final stacks are shown in the figure.

**Figure 2.**
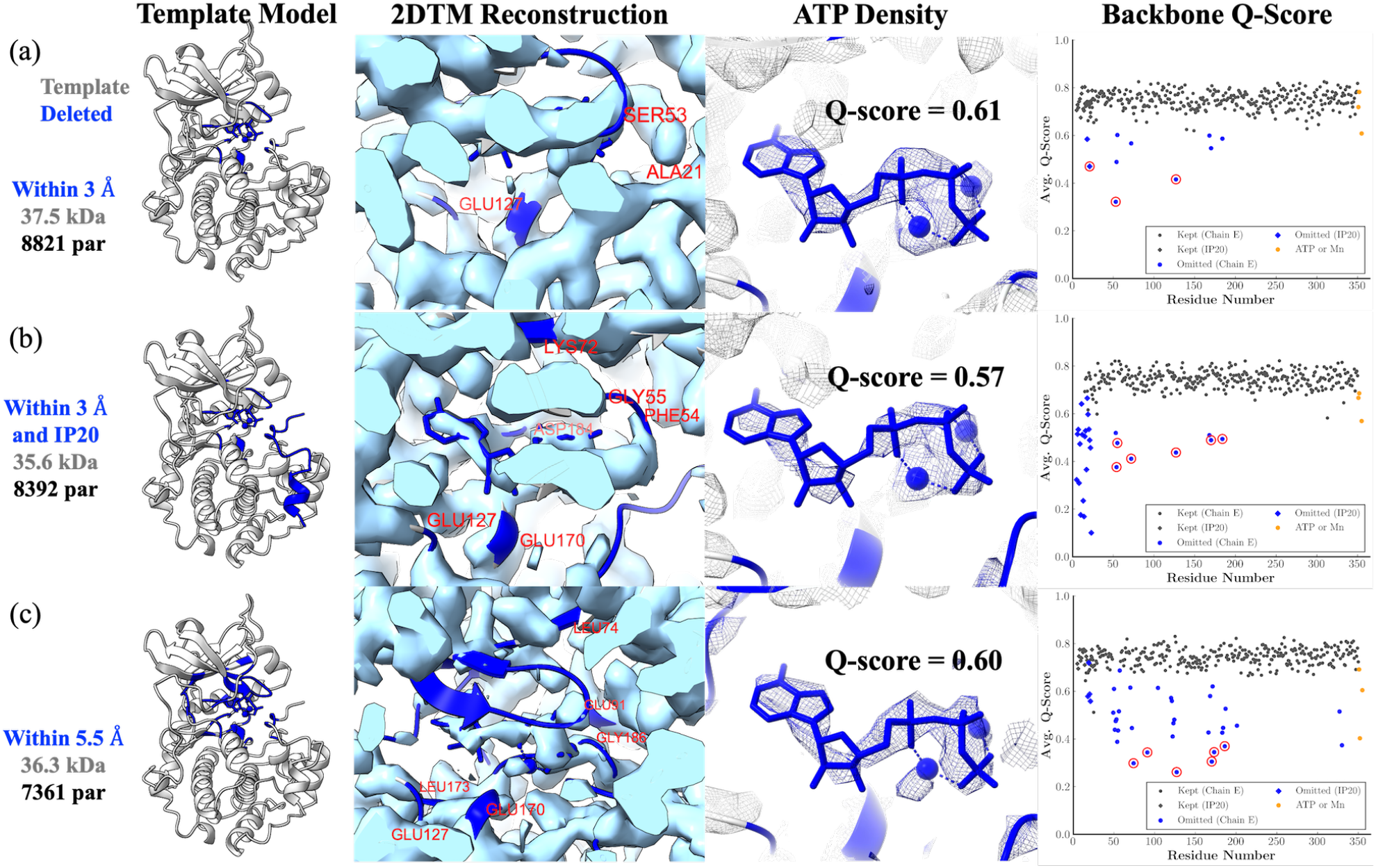
2DTM-based reconstruction of the ATP-binding pocket. Each row shows, from left to right: (1) the atomic model used to generate the template, with residues within a specified radius from ATP deleted (highlighted in blue); (2) the 2DTM reconstruction fitted with the full model, with labeled residues corresponding to those circled in red in the Q-score plot; (3) the recovered ATP density fitted with the full model; and (4) backbone Q-scores of all residues. Omitted residues are shown in blue (circles for chain E, diamonds for IP20); kept residues in gray. ATP and Mn^2+^ are shown in orange. Red circles indicate the omitted residues with the lowest backbone Q-scores, corresponding to the labeled residues in column 2. Q-scores were calculated using the `MapQ` command line tool (34, 35). **(a)** Residues within 3 Å of ATP were deleted. **(b)** IP20 and residues within 3 Å of ATP were deleted. **(c)** Residues within 5.5 Å of ATP were deleted. Map contour *σ* = 5. Map contour σ = 5

In all three cases, the reconstructions showed well-defined ATP and Mn^2+^ densities in the binding pocket. The average Q-scores of ATP in three cases were 0.61, 0.57, and 0.61, respectively. Although some discontinuity was observed at the chosen contour level (*σ* = 5) in Figure 2b, the ATP densities in all reconstructions closely matched the ligand shape in the X-ray model. These results suggest that ATP and Mn^2+^ densities can be robustly recovered from the data, even when the search template lacks these ligands and nearby residues. We also found that nearby protein residues were more affected by template deletion than ATP. When residues within 3 Å of ATP were deleted (Figure 2a), most omitted residues were still recovered with backbone Q-scores above 0.50, except for residues 53 (Ser) and 127 (Glu), which had backbone Q-scores of 0.32 and 0.42, respectively (red circles in Figure 2a). However, when the deletion radius increased to 5.5 Å (Figure 2c), many more omitted residues fell below 0.50, and the densities corresponding to those residues began to show visible discontinuities. This likely was caused by the overall reduction of signal in the template that contributes to the cross-correlation signal. In contrast, ATP density was consistently recovered more robustly than nearby residues. This may be because small angular misalignments disproportionately blur peripheral residues as their displacement scales with distance from the alignment center, whereas the centrally buried ATP density remains relatively unaffected.

Finally, by comparing Figure 2a and b, we observed that deleting IP20 strongly reduced signal at several residues: Phe54, Gly55, Lys72, Glu127, Glu170, and Asp184 all dropped below a backbone Q-score of 0.5 when IP20 was additionally removed, compared with only Ser53 and Glu127 when IP20 was kept. IP20 not only contributes to the overall mass (2.1 kDa) of the template but also sits at the edge of the protein, where it may generate distinct low-resolution features that can facilitate alignment.

Altogether, our experiments demonstrate that a ligand and nearby residues can be deleted to reduce template bias while retaining sufficient signal for their density to be recovered in the reconstruction.

#### Comparison of 2DTM and RELION reconstructions from the same particle stack

To evaluate the quality of particles and poses obtained from 2DTM, we imported the stack of 7,353 particles described in Figure 1 into RELION and performed 3D classification without angular refinement using five classes. As shown in Figure 3a, we combined Class 1-4 into a new stack of 7,197 particles and performed both 3D reconstruction without angular refinement (using the 2DTM-derived orientations directly) and regular (alignment-enabled) 3D auto-refinement in RELION. Each resulting map was then post-processed and low-pass filtered for comparison. As shown in Figure 3b, in both maps, densities for the deleted residues were recovered. Specifically, the densities at ATP and residues 222–227 obtained with the directly imported 2DTM Euler angles were sharper and more continuous than the same region produced with RELION angular refinement. The reconstruction using 2DTM orientations reached 3.1 Å at the FSC=0.143 threshold, compared with 3.7 Å for the RELION auto-refined reconstruction (Figure 3b). We note that neither FSC is a true gold-standard estimate, as both use orientations ultimately derived from the template; however, the comparison between the two processing strategies remains informative. This suggests that the 2DTM-derived orientations are accurate in the omitted regions. We repeated the auto-refinement with different initial low-pass filter values (Figure 3c). The final resolution remained between 3.7 and 4.0 Å, indicating that the starting reference resolution was not the limiting factor in this comparison. We also confirmed that including all five classes yielded 3.7 Å, identical to the reconstruction with Class 5 removed (Figure 3c). A more quantitative assessment of the alignment accuracies attained by 2DTM and RELION requires further work.

**Figure 3.**
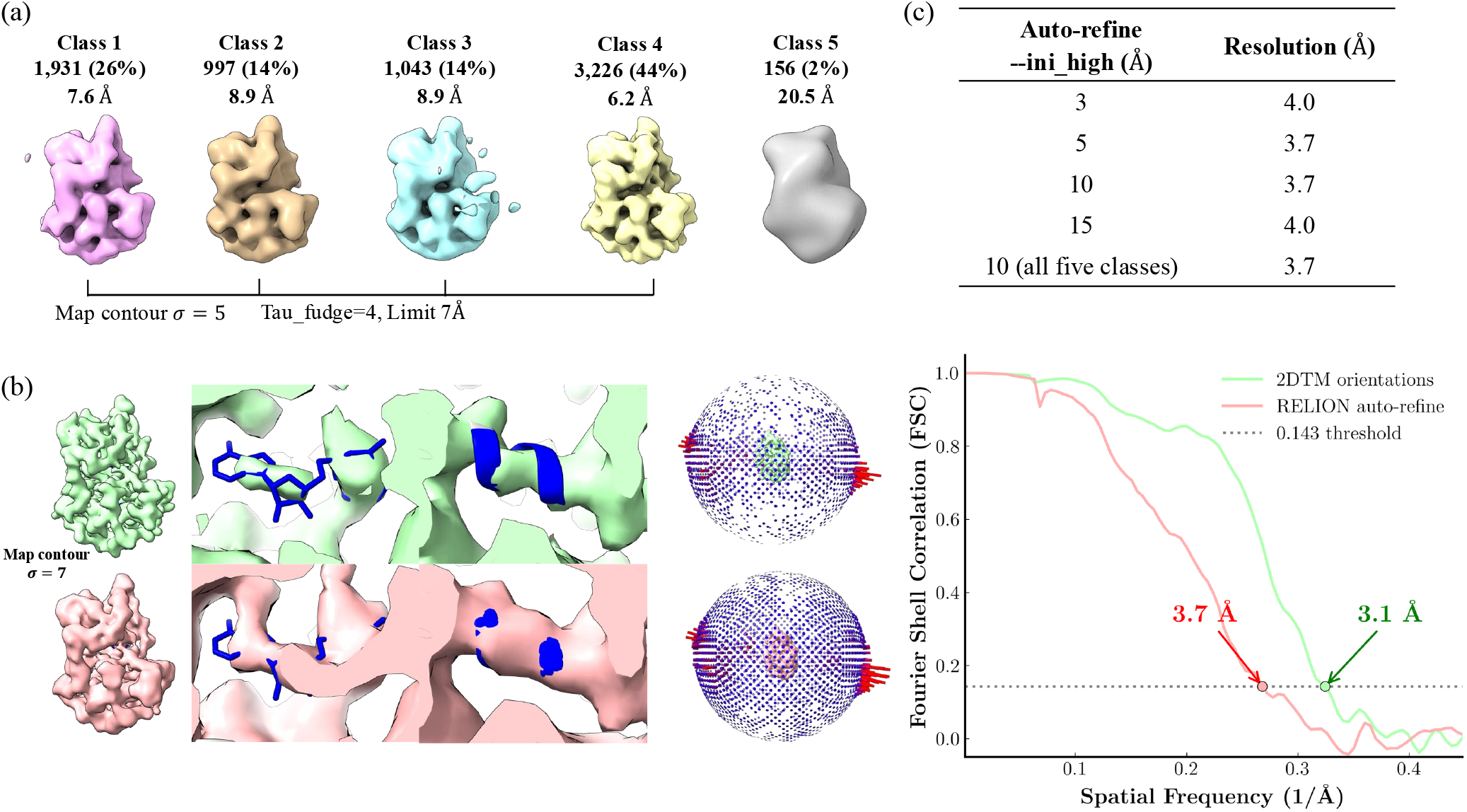
RELION processing of 2DTM-derived particle stack. (a) RELION 3D classification without angular refinement of the 2DTM-derived particle stack from Figure 1, using five classes (Tau_fudge=4, angular limit 7 Å). Classes are colored individually and labeled with particle count, percentage of the dataset, and estimated resolution. Classes 1–4 were merged for 3D reconstruction; Class 5 (156 particles, ∼2%, ∼20.5 Å) was excluded. Map contour *σ* = 5. (b) Comparison of 3D reconstructions using 2DTM orientations directly (top row, green) and RELION auto-refine (bottom row, red). From left to right: full map, zoomed view of the ATP binding pocket and deleted residues 222–227 with the atomic model (blue) overlaid (map contour *σ* = 7), orientation distribution, and Fourier shell correlation (FSC) curves. The 2DTM-orientation reconstruction reaches 3.1 Å and the RELION auto-refined reconstruction 3.7 Å at the FSC=0.143 threshold. In both maps, densities for ATP and the backbone of the deleted residues were recovered, but the 2DTM-derived orientations yielded sharper and more continuous density in the omitted regions. (c) RELION auto-refinement resolution as a function of the initial low-pass filter (`--ini_high`). The final row includes all five classes (i.e., keeping Class 5), yielding 3.7 Å, identical to the reconstruction without Class 5.

### Effective selection strategy enables reconstruction of ATP binding site from around 8k particles

An important difference between our 2DTM approach and the traditional single-particle workflow is the way particles were included in the final 3D reconstruction. In the original publication (30), a final stack of 74,413 particles was used to obtain a reconstruction at ∼ 4.3 Å resolution by gold-standard FSC. The resulting density, however, lacked the expected features at 4 Å, but rather appeared more consistent with a ∼ 6-7 Å map. In particular, the ATP-binding pocket was not resolved in the map (Figure 1a). This particle stack included both untilted and 30^*◦*^-tilted data to reduce preferred orientation. The paper noted that multiple optimizations were attempted, including iterative particle selection based on RELION metadata metrics, but none led to significant improvement.

In our analysis, we implemented a different strategy focused on more stringent particle selection. At the image level, CTF fitting scores from untilted images were computed using CTFFIND5 (36, 37), and only images with scores between 0.05 and 0.2, corresponding to well-fit CTFs, were retained for subsequent 2DTM searches (Figure 1—figure supplement 1a). Examples of micrographs excluded from 2DTM searches are shown in Figure 1—figure supplement 2. At the particle level, we applied two main selection criteria prior to 3D reconstruction:

1. **2DTM-derived statistics:** Due to the small molecular weight of the target, the 2DTM z-score threshold, calculated from the cross-correlations, led to the rejection of most particles and did not yield meaningful detections. Instead, we used the newly developed 2DTM p-value approach to extract particles during the initial processing step (33). Specifically, instead of using the standard first-quadrant p-values, we calculated a three-quadrant p-value to retain particles with low 2DTM z-scores but high 2DTM SNRs (see Methods). We found that a p-value threshold of 8.0 consistently gave us the best reconstruction. Following the initial extraction, as described in the Methods section, we applied additional selection steps based on several 2DTM-derived metrics, including 2DTM SNR, and the pixel-level average and standard deviation of cross-correlations from the angular search.
2. **CTF-based ice thickness:** We excluded images of thicker samples with reduced high-resolution signal, as indicated by their estimated sample thickness from CTF fitting (37). The distribution of mean defocus and thickness of untilted images are shown in Figure 1—figure supplement 1b and c. The mean defocus was 9900 Å. Given that the largest dimension of the protein kinase is ∼ 65 Å, we selected images with estimated thickness between 100 and 800 Å. We found that images with thickness at 300-400 Å contained the most particles based on our criteria (Figure 1—figure supplement 1d and e). Applying these selection criteria drastically reduced the final stack size. For experiments in Figure 1 and Figure 2, around ∼ 8,000 particles were used to generate the reconstruction, only approximately 10% of what was used in the original single-particle pipeline. Despite the order-of-magnitude reduction in data, the resulting 3D map showed a significant improvement at the ATP and IP20 binding sites. However, we note that the global FSC in Figure 1b is not reliable due to the use of the high-resolution template and the resulting template bias (29). Nevertheless, the recovery of density not included in the template confirms that the alignment of the selected particles was sufficiently accurate to generate clear density for the binding sites. Our experiments underline the importance of selecting good particles, rather than maximizing the number of particles selected. Previous studies have pointed out that many particles in the final stack are unnecessary and removing them can improve reconstruction (30, 38). In our experiments, we found that including a larger number of low-quality or misaligned particles, or false positives, may boost the global FSC but blur out weak features such as ligand densities. To test this, we generated a particle stack using a lowered p-value threshold of 7.0. This larger stack (13,669 particles) led to the degradation of signal in the deleted regions, likely due to increased false positives interfering with the true signal (Figure 4). Similarly, applying a 2DTM SNR threshold of 7.5 produced a particle stack of comparable size (8,456 particles) but lower quality, as demonstrated by poor reconstructions in the deleted regions (Figure 4). We also observed preferred orientation as shown by the angular distribution plot of the final stack in Figure 1b. Since we applied very stringent selection, only 2,551 particles were extracted from the tilted images. However, adding these particles did not improve the density at the deleted regions (Figure 4). Although particles from tilted images provide additional angular views, the images often have thicker ice, reducing high-resolution information for accurate alignment. Thus, careful particle curation can enable high-resolution reconstruction of sub-50 kDa complexes with an order-of-magnitude fewer particles.

**Figure 4.**
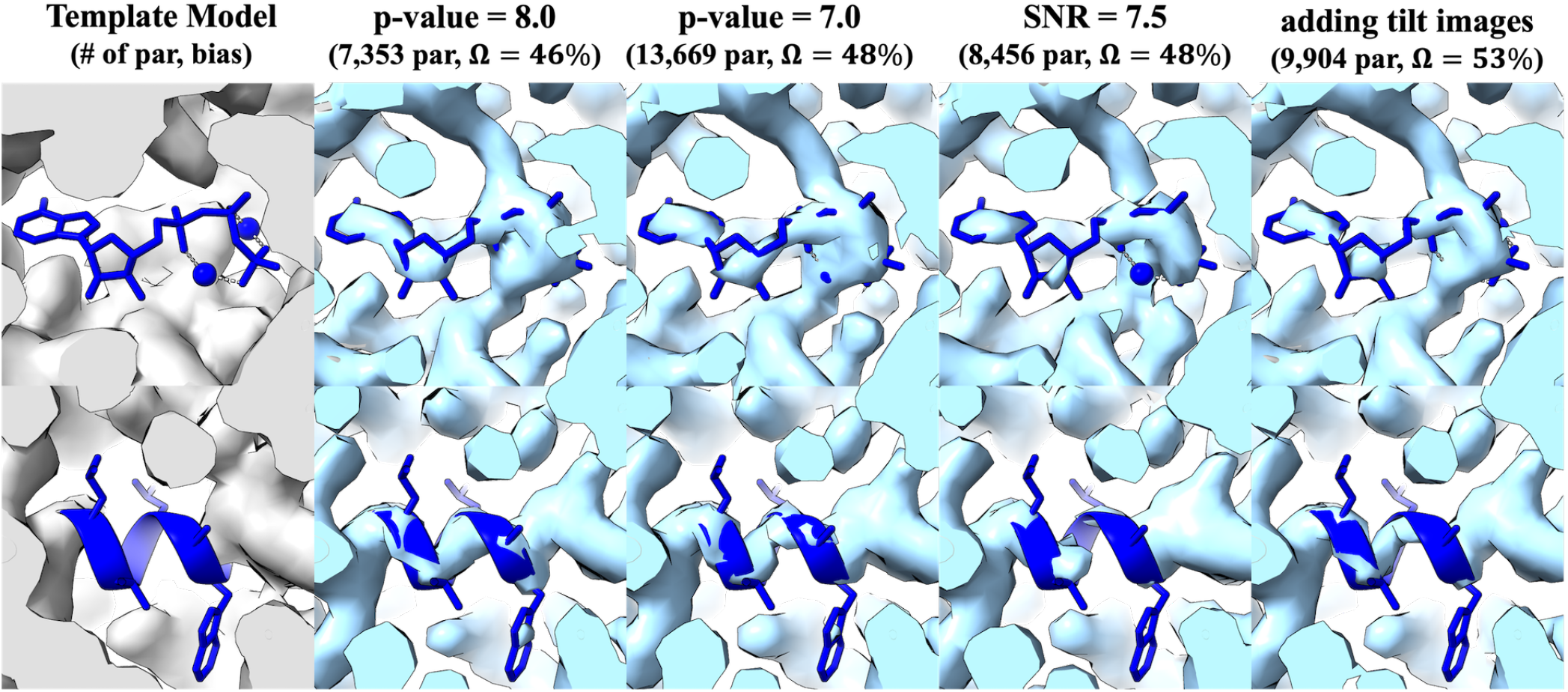
2DTM reconstructions with varying particle selection parameters. The first column shows the template model with the X-ray structure overlaid (deleted atoms in blue). The remaining columns show reconstructions using different selection thresholds (map contour *σ* = 5). The top and bottom rows show zoomed views of the ATP-binding pocket and deleted residues 222–227, respectively. Each column header lists the number of particles and the template bias Ω, defined as Ω = (∑_mask_ *V*_full_ = ∑_mask_ *V*_omit_)*/* ∑_mask_ *V*_full_, where *V*_full_ and *V*_omit_ are reconstructions using orientations and particles derived from independent 2DTM searches with the full and omit templates, respectively, and the sum is restricted to the omission mask derived from the difference between the two templates. Ω = 0 means the omit reconstruction recovers the same density (no bias from the template), while Ω = 1 means all density disappears without the template. Q-scores for ATP and residues 222–227 are reported in Supplementary file 2. Template bias was calculated using the `2DTM_postprocess_tool` Python package, adapted from (29). The omission mask used to compute Ω is shown in Figure 4—figure supplement 1.

### Global density recovery assessed by composite omit maps

To quantify the effect of template bias across different particle selection conditions, we computed a bias metric Ω adapted from (29) (Figure 4). Across the tested conditions, Ω ranged from 46% to 53%, confirming that template bias is present in the reconstruction for features included in the template. To assess whether density could be recovered across the protein when each local region was omitted from the corresponding search template, we performed a composite omit map experiment. We generated 36 omit templates from the X-ray model, each deleting ∼ 10 non-overlapping residues distributed across the protein, including peripheral and surface-exposed regions. For each template, an independent 2DTM search and reconstruction was performed using orientations determined from the corresponding omit-template search. For each reconstruction, density surrounding the omitted residues was extracted and assembled into a composite map (Figure 5), such that each voxel is contributed by a single omit reconstruction (see Methods). This design ensures that the density at each location is derived from a reconstruction in which the corresponding residues were not present in the alignment template. The resulting composite map shows that density can be recovered at distributed locations across the structure, including regions far from the alignment center. Recovery is variable across sites, with some regions exhibiting weaker or fragmented density. This variability is consistent with local differences in structural heterogeneity and residual alignment error.

**Figure 5.**
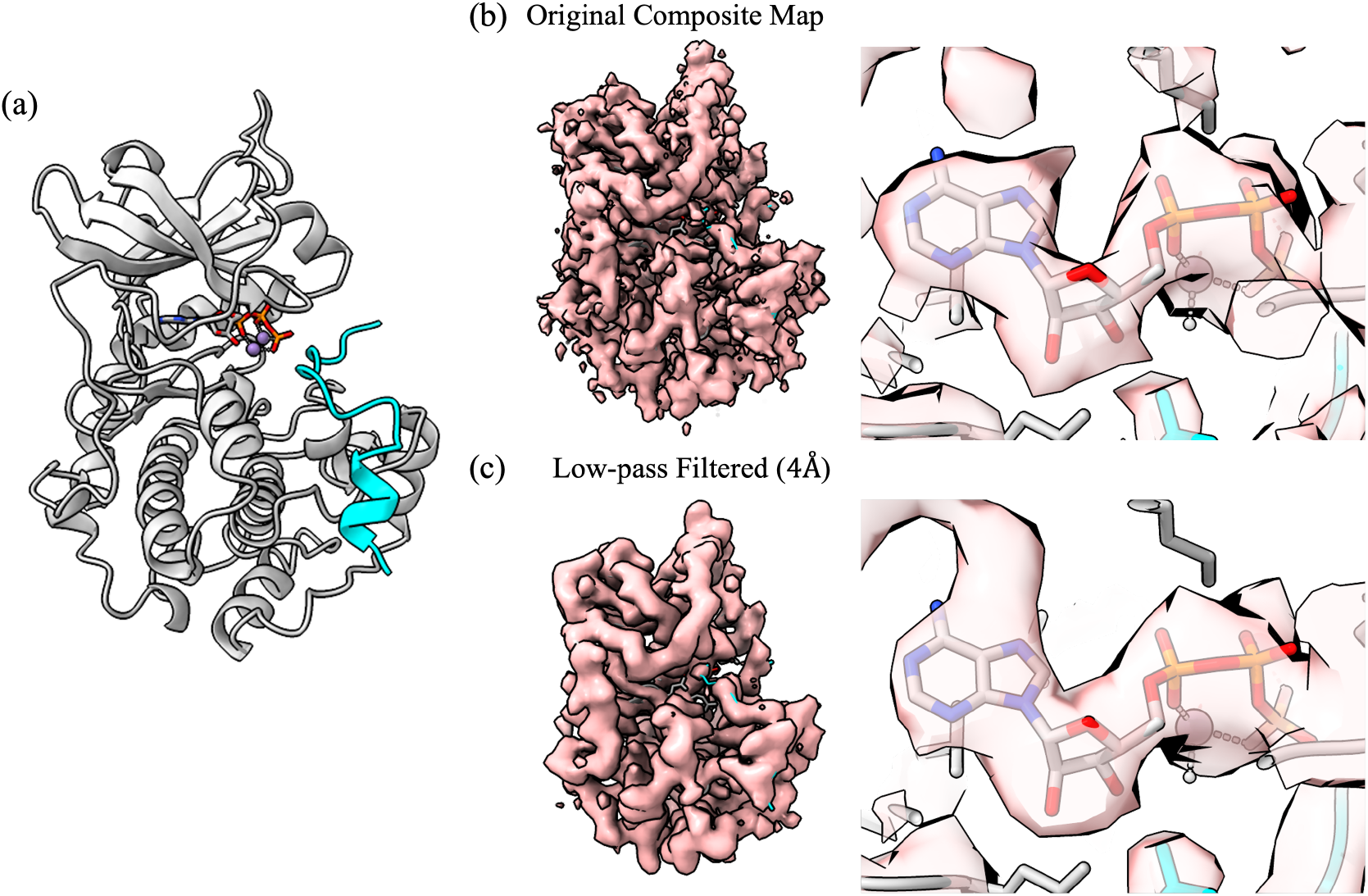
Site-specific composite omit map assembled from 36 partial-deletion reconstructions. Each of the 36 omit templates deletes ∼10 non-overlapping residues. For each template, local density was extracted within 3 Å of the omitted atoms, with neighboring residues (i*±*1) excluded using a 2 Å cutoff on backbone C*α* atoms. Local density patches were assigned uniquely and merged such that each voxel is contributed only by the reconstruction in which the corresponding region was omitted from the template. (a) Atomic model of the protein kinase (PDBID: 1ATP). (b) Left: composite omit map. Right: zoomed view of the ATP-binding pocket. (c) Left: composite omit map low-pass filtered to 4 Å. Right: zoomed view of the ATP-binding pocket. The composite demonstrates that density can be recovered at distributed locations across the protein, including peripheral and surface-exposed regions, although the quality of recovery varies across sites. Map contour *σ* = 5.

To assess the recovered density in Fourier space, we computed a map–model FSC between the composite omit map and a density map simulated from the full 1ATP model (Figure 5— figure supplement 1). The composite map–model FSC crosses 0.5 at 3.5 Å, suggesting an overall resolution of 3.5 Å of the composite map. Because each local region of the composite map was taken from a reconstruction in which the corresponding residues were absent from the search template, this agreement cannot arise from template bias. The resolution estimate for the composite map is lower than the estimate for the reconstruction shown in Figure 1 for several reasons: in the Figure 1 reconstruction, only ATP, Mn^2+^, and residues 222– 227 were omitted, leaving room for template bias, whereas the composite map is assembled from disjoint local omit regions. This stitched construction introduces holes and mask boundaries that affect the FSC curve, in addition to the weaker recovery of locally omitted density.

The composite map–model FSC also showed a negative dip at the lowest spatial frequencies. We hypothesize that this dip is caused by net-negative support-averaged density within the conservative omit masks. Radial binning around the omitted atoms (Supplementary file 3) showed that positive but weakened atom-centered density within ∼ 2 Å is outweighed by the local negative background in the peripheral 2–3 Å shells, making the average density within the omit support negative. We therefore interpret the low-frequency dip as a consequence of support averaging in the stitched omit mask construction rather than as evidence for failed recovery of omitted density.

### Using predicted structures as templates

It is possible that experimental structures are unavailable for the target of interest. We examine whether predicted structures can be used as templates to validate the predictions, or identify novel structures or interactions. We generated a predicted structure for the protein kinase using AlphaFold3 (39). The atomic model includes IP20, ATP, and Mn^2+^ . A high-resolution template was simulated using the same parameters as above using the `simulate` program (32). We show the comparison between the X-ray model and AlphaFold3 model in Figure 6a. Overall, the structures show good agreement, with an RMSD of 0.45 Å across 336 aligned residue pairs. The differences between the AlphaFold3 model and the X-ray structure of protein kinase A are mainly found in the flexible loop regions (e.g., residues 53–55), the surface-exposed side chains, and IP20. In the experiment in Figure 2a, residues within 3 Å of ATP were removed from the X-ray-derived template. Here, the same residues were deleted from the AlphaFold3-derived template, resulting in a remaining molecular weight of 37.3 kDa. Shown in Figure 6b, the densities of ATP in the AlphaFold3-derived reconstruction was slightly worse than that obtained using the X-ray-derived template, highlighting the importance of template accuracy in enabling efficient detection and reconstruction of small complexes. We also observed that both reconstructions resemble closely the search templates, including differences in the side chain densities (Figure 6a). This suggests potential template bias and false positives remained in the particle stack despite the use of stringent selection criteria. Further improvements could involve systematically deleting residues from the predicted model and assessing the impact on the reconstruction, as demonstrated in the composite omit map experiment.

**Figure 6.**
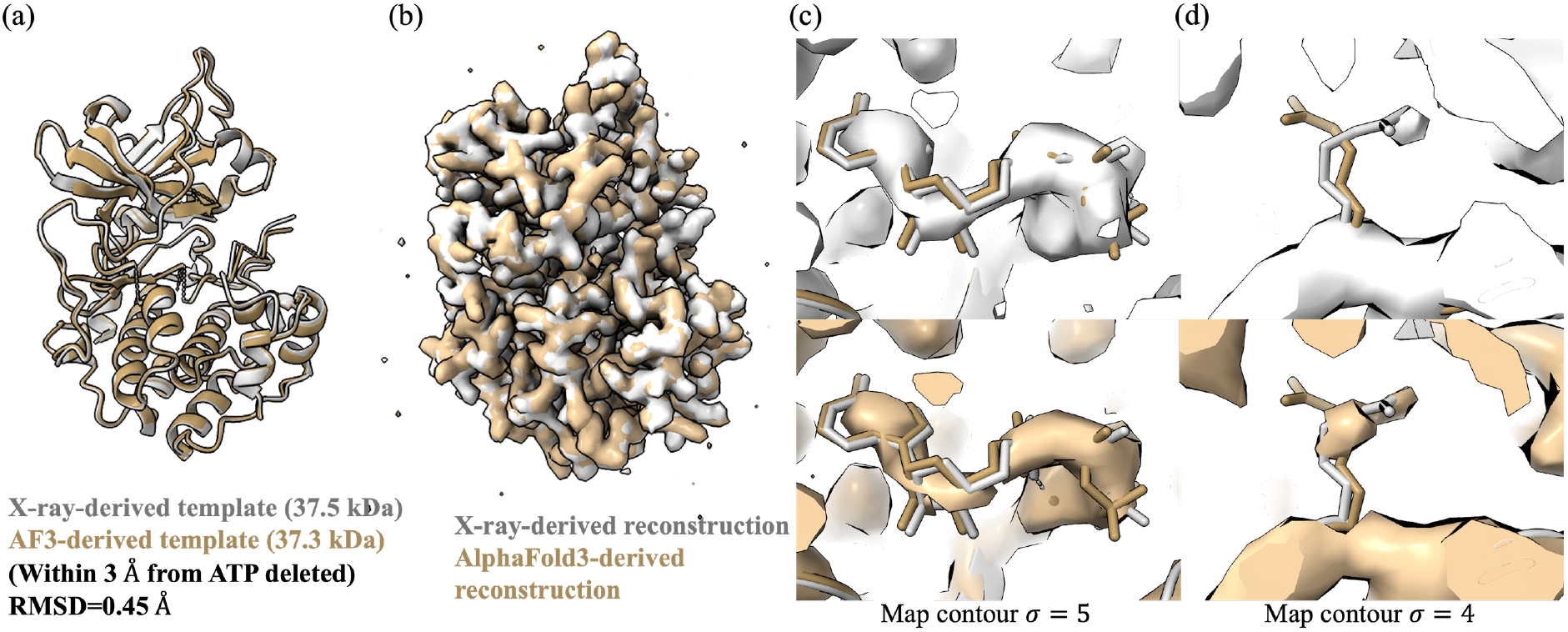
2DTM reconstruction using the AlphaFold model as the search template. (a) Structural comparison between the X-ray model (PDB ID: 1ATP, gray) and the AlphaFold3 predicted model (tan) (39). Residues within 3 Å of ATP were deleted from both templates and not shown. RMSD between the undeleted structures is 0.45 Å. (b) 2DTM-derived maps using the AlphaFold3 template (tan) and the X-ray template (gray). (c) Reconstruction at the ATP-binding site using the X-ray model (top) and the AlphaFold3 model (bottom) as the template (map contour *σ* = 5). (c) Reconstruction at residue 18 (ARG) on IP20 using the X-ray model (top) and the AlphaFold3 model (bottom) as the template (map contour *σ* = 4).

### How small a particle can we study by single-particle cryo-EM

The cross-correlation between the particle image and the reference needs to be larger than the expected cross-correlation between noise and the reference to be alignable (17). This lower molecular weight limit of single-particle cryo-EM was estimated as 38 kDa assuming a total exposure (*N*_*e*_) of 5 e^−^/Å^2^ (16). Later, the estimated limit was lowered to 17 kDa (17). The primary difference between these two predictions lies in the statistical criteria used to assess image visibility: Henderson in (16) required that the intensity of the average Fourier component should be three times the standard deviation of the shot noise, while Glaeser (17) applied the Rose criterion. More recent work has shown that it is now possible to reconstruct proteins below 50 kDa and even smaller nucleic acids, with predictions that the lower molecular weight limit could be extended below 20 kDa (40). Specifically, a high-resolution 3D reconstruction of the 14 kDa hen egg white lysozyme (HEWL) was obtained from a simulated dataset generated with an ideal phase plate (41). More recently, the 40 kDa Aca2–RNA complex was reconstructed to 2.5 Å resolution using Blush regularization, a data-driven denoising prior that improves image alignment for small particles with low signal-to-noise ratios (42). Here, following the rationale of 2DTM, we sought to calculate the lower molecular weight limit for hydrated biological samples for single-particle cryo-EM that takes into account advancements in instrumentation made over the past decades.

During 2DTM, cross-correlation coefficients are calculated between the particle image and 2D projections of the template. *Peaks* in the cross-correlation map indicate regions of high similarities between the image and the template. A signal-to-noise ratio (SNR) can be defined as the maximum correlation observed when aligning an image against a reference, divided by the standard deviation of the correlations from the background. We consider two scenarios: (1) the image contains pure random noise and the corresponding SNR is SNR_*n*_; (2) the image contains real phase contrast from the target and the corresponding SNR is SNR_*s*_. In both scenarios, the SNR is expected to be larger than zero because even in the presence of pure noise, a positive correlation will be obtained after aligning the noise image to a reference. Determining the lower molecular weight limit is equivalent to identifying the intersection at which SNR_*n*_ and SNR_*s*_ are equal. When SNR_*s*_ is smaller than SNR_*n*_, it will be impossible to distinguish signal from noise.

For the first case, cross-correlation coefficients between two pure Gaussian noise images with *N*_*p*_ pixels can be approximated with a Gaussian distribution with zero mean and a variance of 1*/N*_*p*_, where *N*_*p*_ is the number of pixels in the image. In cryo-EM particle alignment, a five-dimensional search is performed for each particle, including two translational parameters (*x* and *y*) and three orientational parameters (*ϕ, θ, ψ*). For each particle, *N*_*s*_ correlation coefficients are calculated to find the correct alignment. *N*_*s*_ is dependent on the size of the particle and the resolution limit of the alignment. A larger particle and higher resolution limit require a more finely sampled search space. The parameter set, Θ_0_ = {*x*_0_, *y*_0_, *ϕ*_0_, *θ*_0_, *ψ*_0_ }, corresponding to the maximum value among *N*_*s*_ correlations, is then used to register the particle’s alignment. We define SNR_*n*_ using the maximum and standard deviation of *N*_*s*_ correlation coefficients. Based on Supporting Information A,

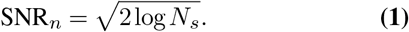

This means the more search locations are evaluated, the higher the likelihood of observing a large correlation value purely by chance.

For the second case, we follow Henderson’s assumption that the particle is roughly spherical and consists of only randomly positioned carbon atoms (16). Cross-correlations are calculated between a *M* -frame summed image and 2D projections from the perfect reference. Similarly, SNR_*s*_ is defined as the ratio between the maximum cross-correlation and the standard deviation of background correlations. Based on Supporting Information B,

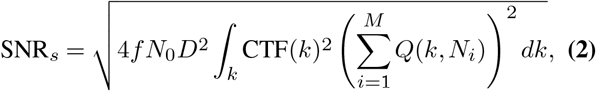

where *Q*(*k, N*_*i*_) is the normalized exposure-weighting transfer function at spatial frequency *k* for cumulative frame exposure *N*_*i*_ (19), *N*_0_ is the exposure per frame, *D* is the particle diameter, and *f* is the fraction of electrons being elastically scattered. Assuming a total exposure of 5 *e*^−^*/*Å^2^ and a single-frame acquisition (16), we derived a simplified form of Eq. 2 by assuming a realistic CTF with multiple oscillations, such that the integral ∫*k* CTF(*k*)2*dk* ≈ 0.5. This leads to

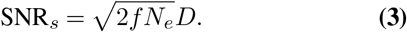

Under these conditions, we estimate a minimum detectable molecular weight of 38.0 kDa. While this result is numerically similar to Henderson’s original estimate (16), the underlying assumptions differ somewhat. (16) assumed ideal imaging conditions without CTF oscillations. In contrast, we assume standard cryo-EM conditions with a realistic CTF and compute the signal and noise terms using different models. In particular, our calculation of the number of correlations calculated (*N*_*s*_) is different. Nevertheless, the agreement in molecular weight limit suggests that this value is a reasonable estimate given the main experimental assumptions (single frame acquisition, 5 *e*^−^*/*Å^2^).

For movies with multiple frames, Eq. 2 can be numerically integrated over spatial frequency range [*k*_min_,*k*_max_] to calculate SNR_*s*_. Eqs. 1 and 2 are then plotted across a range of molecular weights to identify their intersection point, shown in Figure 7. The parameters used for these calculations, along with their corresponding values, are listed in Table 1. Using conventional single-particle analysis conditions with a resolution limit of 2 Å, which can be achieved by collecting images using a pixel size of 1 Å/pixel, assuming a perfect beam (i.e., no envelope function), and using a total exposure of 45 *e*^−^*/*Å^2^, particles as small as 14.8 kDa can be accurately aligned through a full search. If considering the inelastic scattering from ice with a thickness of 30 nm, particles needs to be at least 16.3 kDa to be detected, which is closely consistent with the prediction of 17 kDa in (17). Ideally, if thin ice can be obtained, which is just thick enough to embed the particle, considering the defocus variation across the particle along its diameter, and assuming 10% amplitude contrast, the smallest alignable particle is 14.8 kDa. In practice, such thin ice may be difficult to achieve, leading to an increase in the weight limit. If the low-resolution contrast of the particle allows it to be roughly centered in the x,y plane, we may not need to search the entire area covered by the particle. By constraining the translational search to a 5-by-5 pixel region, the molecular weight limit can be reduced to 11.8 kDa under previously mentioned conditions. Further incorporating a 90^*◦*^ phase plate and using zero defocus lower the limit to 7.4 kDa under constrained search conditions. Previous work has shown that electron diffraction spots fade more slowly at liquid-helium temperatures by a factor between 1.2 and 1.8, compared to those at liquid-nitrogen temperatures (24). Assuming an additional cryo-protection factor of 1.8, particles as small as 5.7 kDa are theoretically alignable by 2DTM.

**Table 1.**
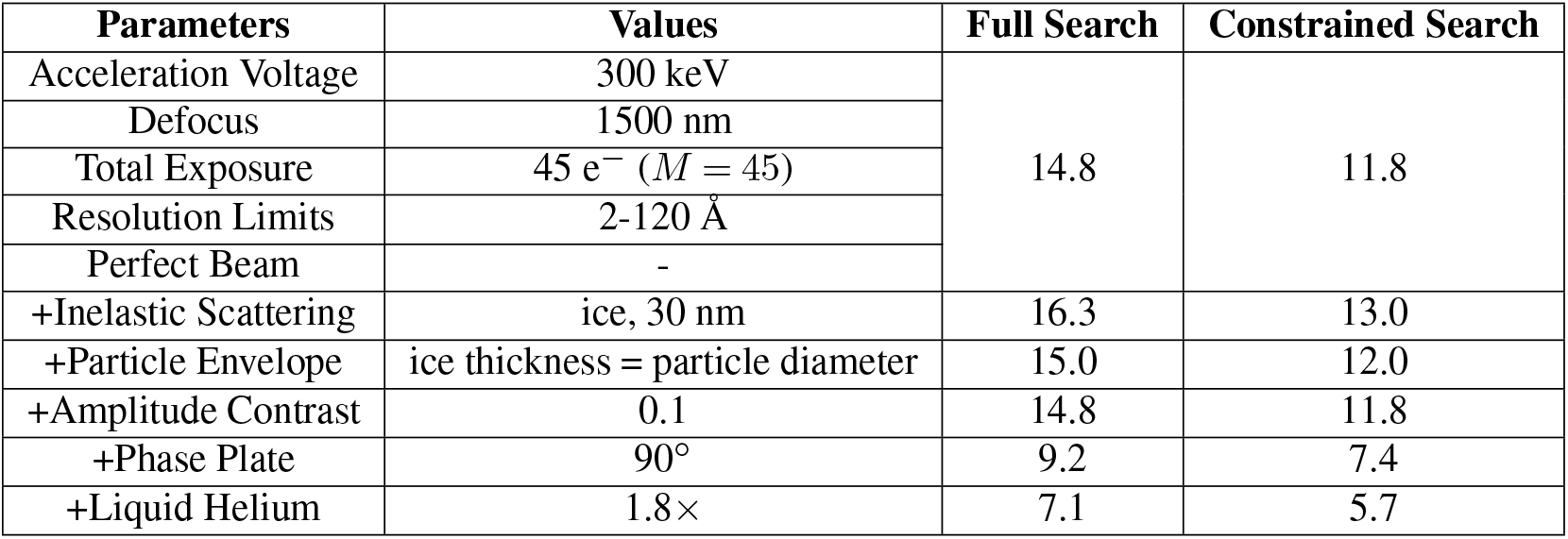
Single-particle cryo-EM lower molecular weight limits under different assumptions. A constrained search restricts the *x* and *y* dimensions to a 5-by-5 pixel window.

**Figure 7.**
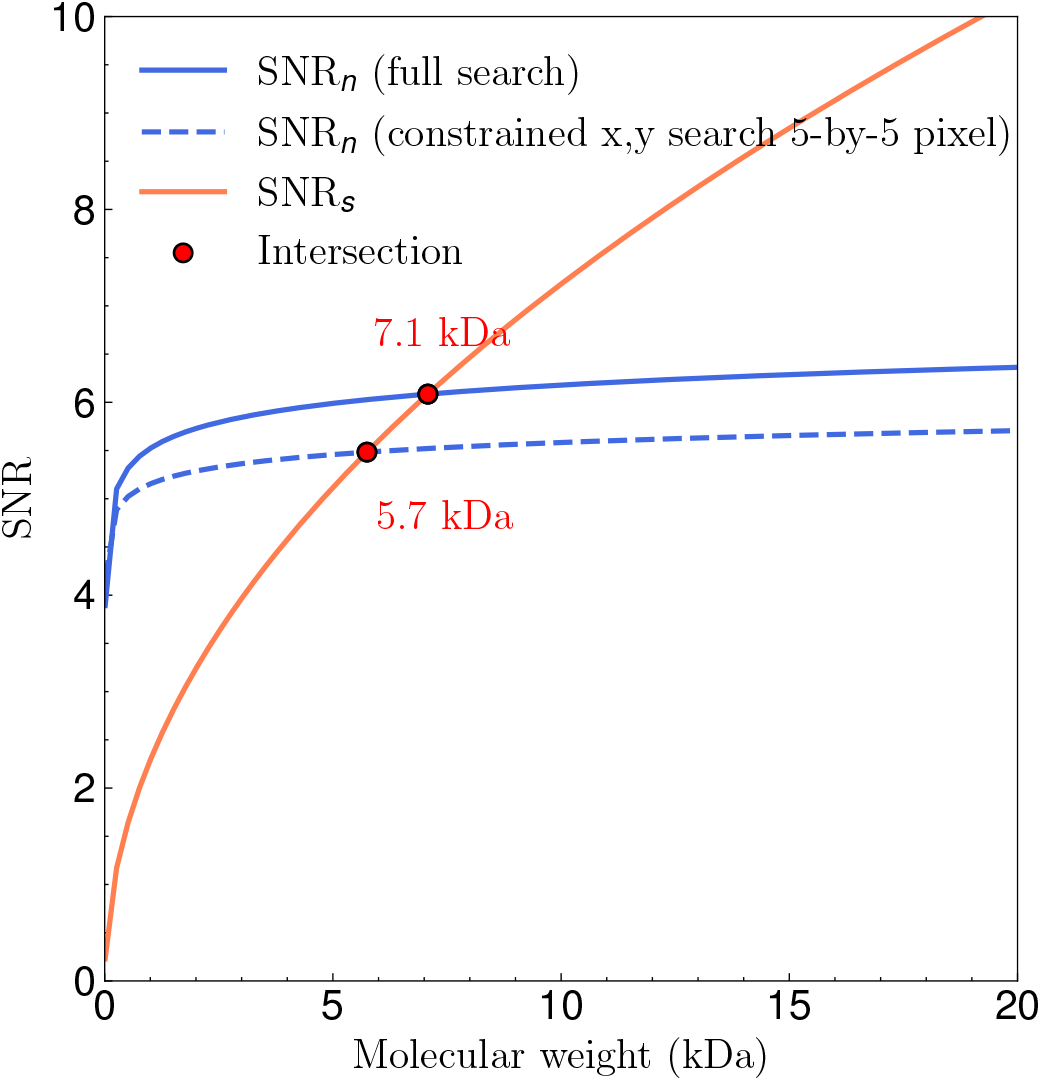
Theoretical lower molecular weight limit. A constrained search restricts the *x* and *y* dimensions to a 5-by-5 pixel window. At the minimal molecular weight, the SNR calculated from alignment noise and phase contrast are equal.

## Discussions

### The reference quality may be a limiting factor of single-particle reconstruction

Images collected in (30) are of high quality, where particles in many images can be visually observed. Our CTF analysis shows that 2,108 images in the untilted dataset exhibited Thon rings extending beyond 4 Å. However, the traditional single-particle processing done in (30) was limited to 4.3 Å resolution, and features at the ATP-binding site were not resolved. In the same work, the**(3)** authors analyzed two other sub-100 kDa complexes, the 82 kDa homodimeric enzyme alcohol dehydrogenase (ADH) and the 64 kDa methemoglobin (metHb), and found that very few particles from many images, rather than many particles from very few images, contributed to the final stack used for reconstruction. This was consistent with our results of selecting a subset of particles for the protein kinase dataset, where most images yielded fewer than 10 particles (Figure 1— figure supplement 1d and e).

So, what is the limiting factor of the single-particle pipeline? In our assessment, we speculate that the images themselves contain sufficient signal to support a reconstruction beyond 4.3 Å resolution. A possible contributing factor is the suboptimal reference used during 3D refinement. In the workflow in (30), an *ab initio* volume generated by cryoSPARC was low-pass filtered to 20 Å and then used as the initial reference for 3D auto-refinement in RELION. Because this starting map lacked high-resolution features, particle alignment may have been less accurate. Once these misalignments were introduced, subsequent refinement iterations may not be able to fully correct them from the local refinement optimum that was created earlier. By contrast, the SAM-IV riboswitch, despite having a comparable molecular weight, has distinct asymmetric features even at low resolution. Combined with the stronger scattering of nucleic acids, this likely facilitated more accurate alignment, allowing the traditional single-particle workflow to achieve a high-resolution reconstruction (43).

### Remaining gaps between experimental and theoretical limits

Despite the progress in cryo-EM, there is still a gap between our predicted lower molecular weight limit in Table. 1 and the smallest template used in our tests (36.3 kDa). Several factors may contribute to this difference:

**Beam-induced motion** particularly affects small particles(44). Two distinct contributions are often considered. The first arises from rapid initial specimen deformation (e.g., cryo-crinkling or buckling of the ice film), which can be substantially reduced through grid design and improved supports (30, 45). The second is a stochastic, pseudo-Brownian motion of particles that accumulates with dose and attenuates high-resolution signal. Based on the analysis of beam-induced motion in amorphous ice and scaling with particle size, this effect is expected to be on the order of a mean-squared displacement of 0.1 Å2 per (e^−^/Å^2^) for a ∼ 10 kDa particle (6). Over a typical total exposure of 40-60 *e*^−^*/*Å2, this corresponds to an accumulated RMS displacement of ∼ 2-2.5 Å, which is sufficient to attenuate medium-to high-resolution signal. Although Bayesian polishing can partially compensate for beam-induced motion, it relies on sufficient particle signal to estimate individual trajectories and assumes spatially smooth motion between nearby particles (46). For very small particles, the per-particle signal may be too weak to reliably support this analysis, and stochastic particle-level motion may therefore remain only partially corrected. The protein kinase dataset was collected using UltrAuFoil grids (30), which reduce initial beam-induced specimen motion; nevertheless, residual motion, particularly pseudo-Brownian motion of small particles, remains a factor limiting the recoverable high-resolution signal in cryo-EM images.

Another factor is the **electron beam energy** used for imaging. The images were collected at 200 keV whereas the theoretical calculation presented here assumes 300 keV. Both elastic and inelastic scattering cross-sections increase at lower voltage. A recent study of sub-200 kDa complexes showed that there is no obvious choice of electron energy for imaging smaller complexes (47). Any gain in image contrast from stronger elastic scattering may be offset by the increasing inelastic scattering (47). However, the lens aberrations are more noticeable at lower energies (48). Detectors are also generally optimized for 300 keV electrons, and practical detector DQE is generally more favorable at 300 keV than at 200 keV. Our calculation assumed a perfect beam whereas the experimental images suffer from spherical and chromatic aberrations. New developments in aberration correction will help obtain atomic-resolution structures of small complexes (49–52).

**Non-ideal experimental conditions** will also widen the gap between theoretical and experimental limits. For example, we assumed the ice is just thick enough to accomodate the particle whereas the images in the protein kinase dataset have varying ice thickness as shown in Figure 1—figure supplement 1c. Additionally, we did not perform a defocus search during 2DTM, instead, we used the average defocus values estimated by CTFFind. Incorporating the defocus search directly into 2DTM may give more accurate estimations in certain cases.

Another contributing factor is the **template generation strategy**. In Figure 1—figure supplement 4, the difference map between the template and the reconstruction for the experiment in Figure 1a was generated using the `diffmap.exe` program (53) with a protein soft mask. Under this masking setup, coherent difference densities are most prominent at the ATP binding site and residues 222– 227, which were deleted from the template. Residual noisy densities in other regions indicate a limitation of the current template generation approach, which may not fully capture the solvent background or accurately model the electrostatic potential of atoms within their molecular environment. As shown by recent work (54) on empirical scattering-factor inference from cryo-EM maps, environment-aware scattering factors that incorporate bonding and charge redistribution effects can improve agreement between model and map. Incorporating such learned scattering factors into template construction would provide a more accurate forward model of the electrostatic potential, potentially reducing spurious residual densities and improving template–map consistency. Additionally, **structural flexibility and conformational heterogeneity** further attenuate high-resolution signal, effectively acting as an additional envelope that reduces the detectable signal relative to the idealized rigid-particle model considered here.

Taken together, these factors explain why the current 2DTM implementation has not yet reached the theoretical lower molecular-weight limit estimated here, and why reliable alignment of proteins much below ∼ 40 kDa remains challenging. Advancements in both microscope optics, detector performance, image processing algorithms, and sample preparation strategies will be essential to close this experimental-theoretical gap and make high-resolution cryo-EM of small complexes a reality.

### Implications for structure-based drug design

For structure-based drug design (SBDD), obtaining a high-resolution structure of the target (and its ligand complex) is a critical step. X-ray crystallography has long been the dominant method for this purpose, but many protein targets are hard to crystallize, especially those that are flexible or membrane-embedded (55). NMR can handle very small proteins or nucleic acids in solution, but typically offers lower resolution information and is limited to proteins below 25 kDa, otherwise spectra become too complex to interpret without isotope labeling and extra processing (8, 9). Cryo-EM is emerging as a powerful alternative and complementary method and enables structural and functional studies under conditions more closely resembling the native cellular context. One of the major caveat for cryo-EM, however, is the reconstruction of sub-50 kDa complexes.

The 2DTM-based single-particle alignment and reconstruction workflow we proposed here **simplifies** the conventional single-particle pipeline by foregoing iterative rounds of 2D classification, *ab initio* modeling, 3D classification and refinement. 2DTM directly returns particles with their *x, y* position, Euler angles and defocus in *one pass*. Aside from the computational cost of the search itself, the workflow is trivial compared to a single-particle pipeline. For a typical single-particle dataset of ∼ 2,000 micrographs (5k *×* 4k pixels), a 2DTM search without defocus refinement completes in approximately one day on 64 NVIDIA A6000 GPUs. Once particles are located with their orientations and positions, a single 3D reconstruction is sufficient without further refinement, eliminating the iterative 2D classification, *ab initio* modeling, 3D classification and refinement steps of a conventional pipeline.

Another motivation for 2DTM is overcoming the **size limit** of cryo-EM. Conventional single-particle workflows have struggled to achieve good reconstructions for particles below 50 kDa, where low contrast makes alignment unreliable with an imperfect reference. To date, only a handful of isolated sub-100 kDa structures have been solved at *<*4 Å resolution. Notable examples include the 52 kDa streptavidin tetramer resolved to 3.2 Å using a Volta phase plate and C_s_ corrector (56), and to 2.6 Å using graphene grids (57); the 64 kDa methemoglobin at ∼ 2.8 Å resolution (30); the 39 kDa SAM-IV riboswitch at 3.7 Å resolution (43), and the previously mentioned 40 kDa Aca2–RNA complex at 2.5 Å (42). In a recent study reporting the 3D reconstruction of the protein kinase also used in the present study, the authors achieved near-atomic resolution with a carefully tuned *ab initio* procedure, using a narrow band of spatial frequencies that was centered at 5 Å and incremented in small steps between reconstruction iterations to 2.3 Å resolution (58). These complementary findings highlight the importance of high-resolution information for small particle alignment. In our case, the 2DTM-based reconstruction method improves alignment by explicitly utilizing the high-resolution information from a perfect reference, enabling robust recovery with far fewer particles. Our theoretical estimation of the lower molecular weight limit further highlights that for perfect images and a perfect reference, complexes as small as 5.7 kDa can be accurately aligned when using liquid helium cooling and a phase plate. The workflow we describe here also has the potential to operate as a **screening platform**, enabling structure-guided optimization of small-molecule ligands even for targets below the traditional cryo-EM size limit. High-resolution cryo-EM is already being applied to drug targets that resist crystallization: for example, the high-resolution (up to 1.8 Å) maps of human CDK-activating kinase bound to 15 different inhibitors revealed detailed inhibitor interactions and water networks in the active sites (59). In another case, cryo-EM captured a novel allosteric mechanism for protein inhibition of the human ATP-citrate lyase that enhances the target’s “druggability” (60). These studies show that resolving the ligand-bound structures can directly guide design of novel therapeutics. Our workflow can broaden this approach to smaller drug targets. In principle, one could incubate a sub-50 kDa target with various inhibitors then apply 2DTM using the apo structure as the template, which can be determined from *in vitro* experiment or AlphaFold predictions. The ligand-bound complex can be located, aligned and reconstructed using 2DTM. 2DTM thus offers a structure-based assay: binding of each inhibitor produces a distinct density feature at the binding site, streamlining hit validation.

In summary, by overcoming the ∼ 50 kDa barrier, 2DTM broadens the possibilities for structural studies of many previously inaccessible drug targets that have remained difficult to reconstruct, with the ultimate goal of integrating cryo-EM into high-throughput SBDD.

### Toward data-driven refinement of AlphaFold3 models via 2DTM

Our results demonstrate that AlphaFold3-predicted structures can be used directly as templates in 2DTM searches to pick particles and reconstruct high-resolution maps, even in the absence of any experimentally determined model. This validates the use of AlphaFold3 as a starting point for structure determination. While the predicted structure may deviate from the true conformation, as shown in Figure 6a, the resulting cryo-EM map provides information that can guide further refinement of the AlphaFold3 model. This refined model could then be used as an improved template in a second round of particle picking and reconstruction, allowing recovery of more accurate particles and a higher-quality map. In Figure 6d, we show a deleted residue and the reconstructed densities obtained using both the X-ray template and the AlphaFold3 template. Although the densities are weak and discontinuous, both maps are consistent with the side chain conformation observed in the X-ray model. While we have not yet tested this type of refinement, these preliminary results suggest that this data-driven approach could progressively improve both the structural model and the final reconstruction, starting entirely from prediction. We have demonstrated residue-level omission validation using the X-ray template via a composite omit map (Figure 5); however, extending this approach to AlphaFold3-based templates remains computationally expensive and we consider this an important direction for future work.

A more systematic benchmark of 2DTM against recent single-particle methods for small particles, such as Blush regularization (42) and HR-HAIR (58), will be an important next step. Such comparisons should consider the size of the particle stack, the molecular mass of the particle, and the fidelity of the template forward model. In the ideal limit of an accurate template and forward model, 2DTM should provide a perfect prior for particle detection and pose determination. In practice, current templates remain imperfect approximations to the experimental signal because of solvent-boundary effects, atomic scattering-factor accuracy, bonding and charge redistribution, and conformational mismatch. Together with improved template generation models, such benchmarks will help define the most appropriate use cases for 2DTM, including high-resolution pose initialization, particle-stack enrichment, and ligand-focused screening.

## Methods

### Cryo-EM data set and image processing

Unaligned movies of the protein kinase were downloaded from EMPIAR-10252 (30). The dataset contains 4,809 images, among which 2,488 are from untilted samples and 2,321 are collected at 30^*◦*^ tilt. Motion correction and exposure-weighting were performed using the MotionCor2 program (61). We kept the same procedure as the original publication by using 5 *×* 5 tiled frames with a B-factor of 250 Å^2^ and a binning factor of 2. We used the exposure weighted summed frames for CTF fitting using CTFFind5 (37) with a box size of 512 and a resolution range of 4-30 Å. We selected images with CTF fitting scores between 0.05 and 0.2 for the untilted dataset and 0.04-0.13 for the tilted dataset. Images that were excluded are shown in Figure 1—figure supplement 2 for comparison.

### 2DTM search

After selecting good micrographs based on their CTF fit, 2,314 images of untilted and 2,252 of tilted samples with a pixel size of 1.117 Å/pixel were used for 2DTM. For the experiment in Figure 1, ATP, Mn^2+^, and residues 222–227 were deleted from the X-ray model (PDBID: 1ATP) before template simulation. For the experiment in Figure 2, residues within a radius of 3 Å and 5.5 Å from ATP were deleted from the X-ray model. IP20 was also deleted for the test in Figure 2b. Modified atomic coordinates were generated using UCSF ChimeraX (62). High-resolution 3D templates were then generated from the modified models using program `simulate` in *cis*TEM (32). A uniform B-factor of 30 Å^2^ was applied to all atoms. 2DTM searches were done using an angular search step of 2.5^*◦*^ for out-of-plane angles and 1.5^*◦*^ for in-plane angle for all tests with no defocus search.

### Particle extraction and selection

To streamline post-processing of 2DTM, we implemented a dedicated Python toolkit `2DTM_postprocess_tool`. The module contains two command line tools:

`1. extract-particles`. Input arguments are (i) the *cis*TEM 2DTM SQLite project database, (ii) the IDs of the search and associated CTF-estimation jobs, and (iii) the image pixel size. The program extracts candidate particles in each image in the specified job using one of the three 2DTM metrics: SNR, z-score, or p-value.

For the p-value, the user may choose first-quadrant (1Q) or three-quadrant (3Q) definitions (33): after probit transformation of z-score and SNR, 1Q keeps candidates where both probit-zscore and probit-SNR are greater than 0, while 3Q keeps candidates where at least one of the two values is greater than 0. In the protein kinase example, we first located local maxima in 2DTM SNR maps (exclusion radius = 10 pixels; micrograph border mask = 92 pixels to avoid truncated particles), then calculated the three-quadrant p-values and stored particles with p-values larger or equal to 8.0.

`2. filter-particles`. This function provides secondary quality selection metrics based on the CTF fitting quality, sample thickness, and particle-level statistics from the 2DTM angular search (mean and standard deviation of per-pixel cross-correlations across sampled orientations). For the present study we required a SNR*>* 6.0, an ice thickness between 100 and 800 Å, and a CTF fitting score between 0.05-0.2 for untilted images (or 0.04-0.13 for tilted). Particles whose angular search mean cross-correlation was negative or standard deviation of cross-correlations exceeded 1.1 were also discarded.

Both steps output a star file that can be used for the following 3D reconstruction.

### 3D reconstruction

Particle stacks processed by `filter-particles` and alignment parameters were imported into *cis*TEM as a refinement package for single-particle processing. A 3D reconstruction was generated using the *cis*TEM program `reconstruct3d`. UCSF ChimeraX was used for visualizing the final reconstructions.

### RELION processing

Particles selected by 2DTM were subjected to 3D classification in RELION using a tau fudge factor of 4 and an E-step resolution limit of 7 Å, resulting in five classes. The best-resolved classes were subsequently used for either (i) 3D reconstruction without angular refinement, using the 2DTM-derived orientations directly, or (ii) 3D auto-refinement with alignment, using C1 symmetry and a 3.7^*◦*^ angular sampling step with the corresponding 10 Å low-pass filtered template as reference (`--ini_high` 10 Å). A soft mask was then generated from the resulting map by applying a 15 Å low-pass filter and using a soft-edge of 6 pixels. The final map was produced by post-processing with B-factor sharpening and low-pass filtering. To test the effect of the initial reference resolution, auto-refinement was additionally repeated with `--ini_high` values of 3, 5, and 15 Å. The all-five-class reconstruction was performed using `--ini_high` 10 Å with the same masking and postprocessing protocol.

### Template bias quantification

To quantify template bias in the omitted region, we computed the metric *V*_full_ and *V*_omit_ are 3D reconstructions generated using Ω = (∑_mask_ *V*full − ∑_mask_ *V*omit)*/* ∑_mask_ *V*full, where orientations and particles derived from independent 2DTM searches with the full and omit templates, respectively (29). The omission mask was derived by subtracting the omit template from the full template and thresholding at one-fifth of the mean of the top-100 difference voxels, yielding 959 masked voxels concentrated at the ATP-binding site and residues 222–227 (Figure 4—figure supplement 1). Ω = 0 indicates no template bias (the omit reconstruction recovers identical density), while Ω = 1 indicates full bias (all density vanishes when not included in the template). Bias was computed using the `measure-template-bias` function in the `2DTM_postprocess_tool` Python package (https://github.com/kekexinz/2DTM_postprocess_tool), also available in the official *cis*TEM repository (https://github.com/timothygrant80/cisTEM) (29, 33).

### Composite omit map construction

To avoid template bias and test spatial generalization of density recovery, we constructed a composite omit map from multiple independent reconstructions. Starting from the X-ray model (PDBID: 1ATP), we generated 36 omit templates, each deleting a set of ∼10 non-overlapping residues distributed across the structure.

For each omit template, a separate 2DTM search was performed, and particle orientations obtained from that search were used to compute a corresponding 3D reconstruction.

For each omit reconstruction *k*, a binary mask *M*_*k*_(**x**) was defined to isolate density associated with the omitted residues. Voxels within 3 Å of any atom in the omitted residues were included, while voxels within 2 Å of backbone C*α* atoms of adjacent residues (i *±* 1 in sequence) were excluded to reduce contamination from neighboring template-included regions. To ensure that each voxel in the final composite map is contributed by at most one reconstruction, overlapping mask regions were resolved deterministically. A support map was first computed to identify voxels covered by multiple masks. For voxels with multiple assignments, the contributing reconstruction was selected as the one whose nearest omitted atom is closest to the voxel.

Each binary mask was then converted to a soft mask by applying a cosine-edge taper using a distance transform, with a taper width of two voxels, ensuring smooth transitions without expanding the support beyond the assigned region. The final composite map was constructed as

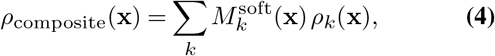

where *ρ*_*k*_(**x**) is the reconstruction corresponding to omit template *k*, and 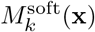 is the tapered mask. By construction, each voxel in the composite map is derived from a single omit reconstruction.

Uncorrected half-map FSCs of the omit reconstructions were computed with *cis*TEM `calculate_fsc`. For each of the 36 partial-deletion reconstructions, the FSC was computed between its two native half-maps (Figure 5—figure supplement 2). For the composite, two half-maps were assembled from the 36 pairs with the same voxel-assignment masks 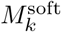 applied to both halves. Because this imposes identical voxel provenance in both halves, the resulting FSC is inflated, as shown by a phase-randomization control (63) in which the parent half-maps’ phases beyond 4 Å were independently randomized and reassembled with the same masks (Figure 5—figure supplement 3). The composite half-map FSC is therefore not a reliable resolution estimate.

## Code availability

The `2DTM_postprocess_tool` Python package, which includes the `extract-particles`, `filter-particles`, and `measure-template-bias` command line tools used in this work, is available at https://github.com/kekexinz/2DTM_postprocess_tool and in the official *cis*TEM repository at https://github.com/timothygrant80/cisTEM.

## ACKNOWLEDGEMENTS

We thank the members of the Grigorieff lab for the fruitful discussion of this work. We are especially grateful to Dongjie Zhu for sharing and testing his new methods and for many insightful conversations.

## Supplementary Note 1: Estimating the lower molecular weight limit

### A. SNR from alignment noise

For two independent Gaussian noise images of *N*_*p*_ pixels, the normalized cross-correlation, *r*_*n*_, is zero on average. The distribution of *r*_*n*_ when *N*_*p*_ is large can be approximated by a Gaussian distribution

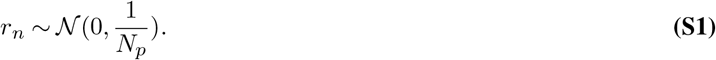

In cryo-EM particle alignment, normalized cross-correlations are computed between a noisy particle image and a set of clean 2D projections generated at sampled orientations, in order to determine the best-matching position (*x, y*) and orientation. For each particle image, *N*_*s*_ cross-correlations are evaluated, and the alignment is assigned based on the position and orientation corresponding to the maximum cross-correlation. This process is equivalent to drawing the maximal value from *N*_*s*_ Gaussian distributed random variables with zero mean and variance of 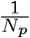. The upper bound of the expectation of this maximum is (65)

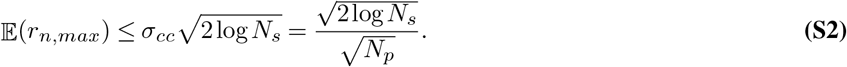

We define the signal-to-noise ratio (SNR) of alignment noise as the number of standard deviations (SDs) by which the maximum cross-correlation exceeds the SD of cross-correlations computed across a pure noise image:

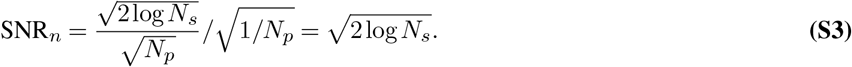

Assuming that the high-resolution limit for alignment is *d* = 1*/k*_max_ (Å), where *k*_max_ is the maximum spatial frequency (Å^−1^). The ideal pixel size is then *p* = *d/*2 (Å/pixel). For a particle of diameter *D* (Å), a five-dimensional search includes the following components:

1. in-plane rotations: 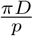
2. Out-of-plane viewing directions:

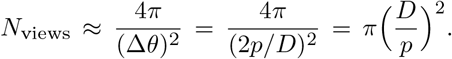
3. *x, y* shifts: 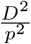

The total number of correlations calculated during a five-dimensional search is then

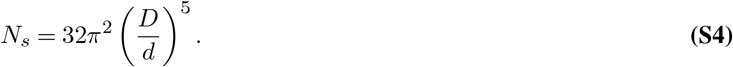

### B. SNR from phase contrast

#### Fraction of electrons being elastically scattered up to a resolution limit

When the image contains phase contrast, the SNR is defined as the number of SDs by which the normalized cross-correlation exceeds the background SD, consistent with the definition used in the 2DTM implementation (26).

We assume that the particle consists of only randomly positioned carbon atoms. The electron atomic scattering factor of carbon can be approximated as a sum of (normally) five Gaussians (66, 67):

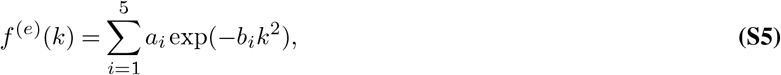

where *a*_*i*_ and *b*_*i*_ are the fitting parameters up to 6 Å^−1^ (68), and *k* is the spatial frequency (Å^−1^). The differential scattering cross-section is:

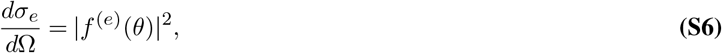

where *θ* is the scattering angle and

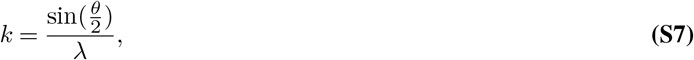

where *λ* is the electron wavelength. The differential cross-section for a single, isolated atom is related to *θ* by:

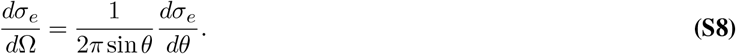

We now integrate to calculate the total scattering cross-section:

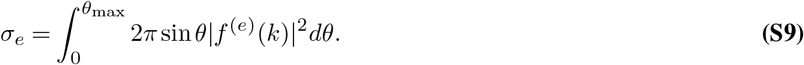

Assuming protein density *ρ* ≈ 0.8 Da/Å^3^, the number of carbon atoms equivalent to a spherical protein of diameter *D* (Å) is (16):

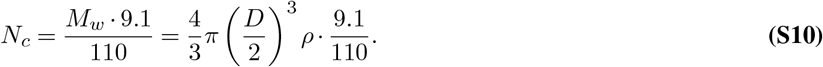

The fraction of electrons being elastically scattered by the particle is:

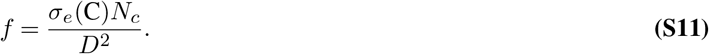

The exit wave function at distance *z* below the specimen is (69)

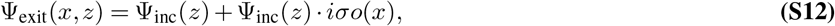

where *o*(*x*) is the projected potential of the molecule. We can now relate the molecular weight of the particle to the Fourier component of the image:

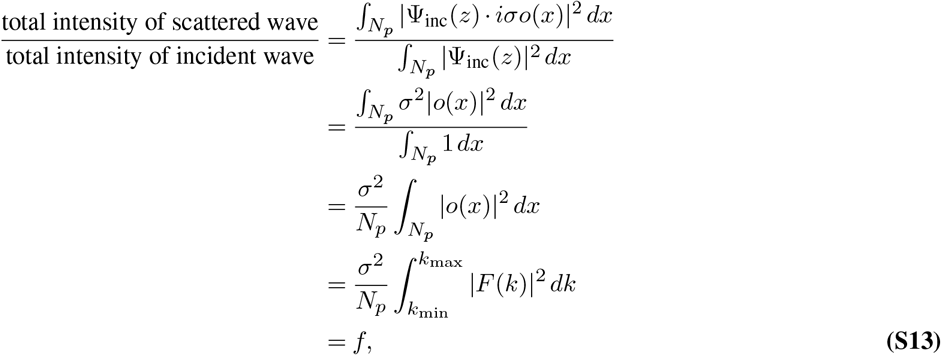

where *F* (*k*) denotes the 2D Fourier transform of the projected Coulomb potential *o*(*x*) of the particle.

#### Image formation model

The wave function in Fourier space after lens aberration is (6, 69):

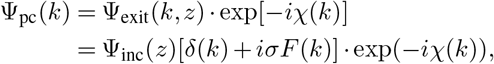

in which the lens aberration function (70) is

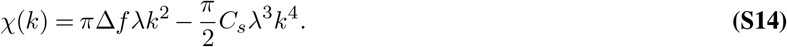

Here, Δ*f* is the defocus (positive for underfocus) and *C*_*s*_ is the spherical aberration. The contrast transfer function (CTF) is defined as

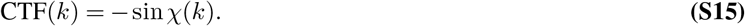

The Fourier transform of the linearized intensity is (6, 69):

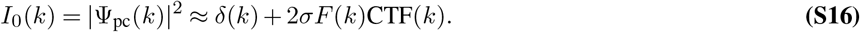

Eq. S16 is the *linear model of image formation* in cryo-EM.

Given the per-frame per-unit area exposure frame is 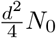 where *N*_0_ is the exposure per-frame, the observed noisy image of a single

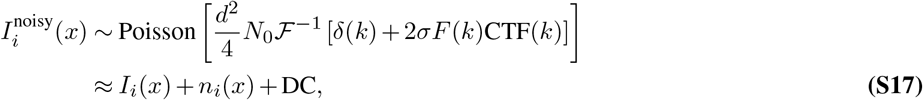

where

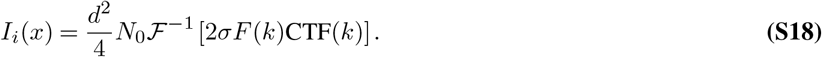

Equivalently, in Fourier space

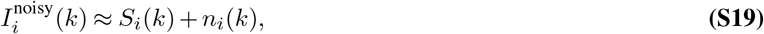

where

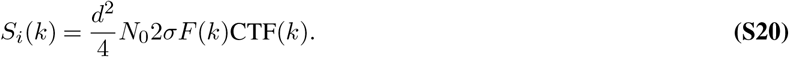

And the noise term *n*_*i*_(*x*) is additive white Gaussian noise:

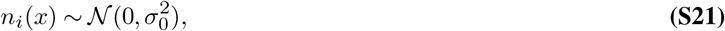

where 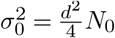. We will ignore the contribution of the DC term in the cross-correlation.

#### Matching with a perfect reference

We now define an SNR value as the expected value of cross-correlations generated from phase contrast, divided by the SD of the correlations from noise

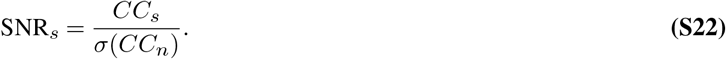

For an image summed over *M* frames, Eq. S19 is updated with an exposure filter *Q*(*k, N*_*i*_) calculated from (19). Here, *Q*(*k, N*_*i*_) denotes the normalized exposure-weighting transfer function at spatial frequency *k* for frame dose *N*_*i*_ (*N*_*i*_ = *i N*_0_). The unnormalized exposure-dependent weight is

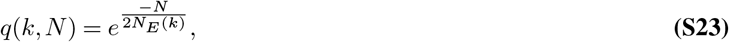

and the normalized filter is

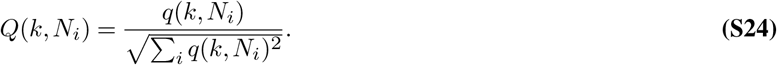

The processed image is then

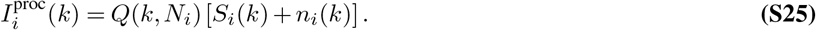

It follows that

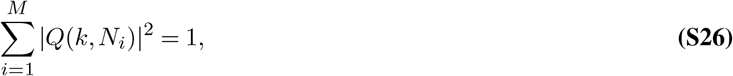

ensuring the noise in the summed frame remains “white” with variance *σ*^2^. Additional instrument-dependent attenuation factors, such as detector DQE and temporal or spatial coherence envelope functions, were not explicitly included. These effects act as additional frequency-dependent weighting terms in Fourier space and could be incorporated into the same filter without changing the structure of the derivation.

The summed image:

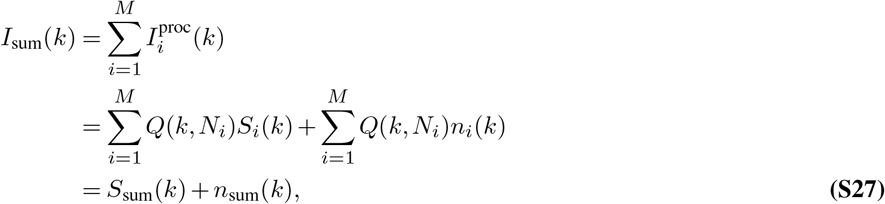

where

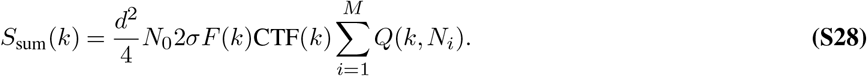

Using a perfect reference that is also exposure filtered

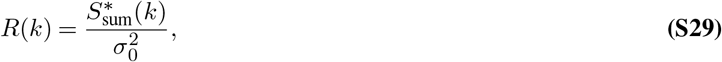

the expected CC from the signal is

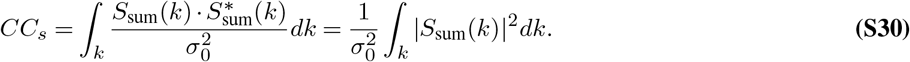

For noise, the mean CC is zero and the variance is:

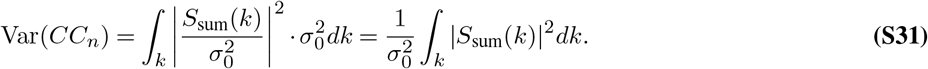

Thus, the SNR from phase contrast, assuming Wilson statistics (71) (flat spectrum of randomly positioned carbon atoms), is:

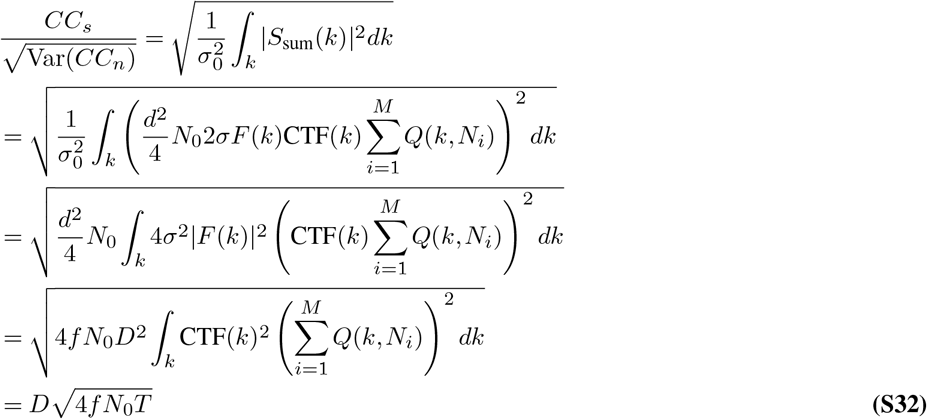

where the integral *T* can be calculated numerically.

### C. Interpretation when SNR_*n*_ **= SNR**_*s*_

Eq. 1 describes an *expected* alignment-noise level from the maximum over *N*_*s*_ tested hypotheses, where *N*_*s*_ includes in-plane rotation, out-of-plane viewing directions, and *x, y* shifts for a per-particle 5D search. Therefore, SNR_*n*_ = SNR_*s*_ should be interpreted as a threshold point in expectation, not a deterministic boundary for each particle. When SNR_*s*_ is only slightly above SNR_*n*_, correct alignments are favored on average and real density can accumulate with sufficiently large particle numbers, but residual pose errors still attenuate high-resolution amplitudes (effectively a larger positive B-factor). In this regime, sharpening (negative-B correction) can improve visibility after averaging, but cannot recover information lost by misalignment.

### D. Advanced assumptions

#### Inelastic scattering

We further add a correction for inelastic scattering (assuming the use of an energy filter to remove inelastically scattered electrons) using:

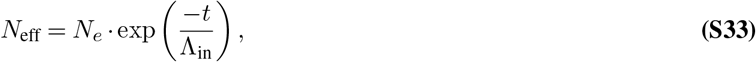

where *t* is the sample thickness and Λ_in_ is the inelastic scattering mean free path of the solution. The inelastic mean free path depends on the incident electron energy and is related to the scattering cross-section as:

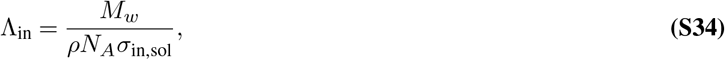

where *σ*_in,sol_ is the weighted inelastic scattering cross-section calculated by individual atom’s inelastic scattering cross-sections (72, 73):

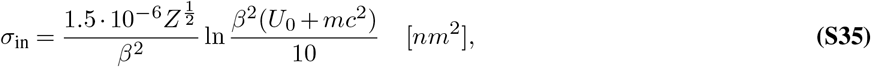

where *Z* is atomic number, *β* the ratio between velocity of the electron and light, *U*_0_ the incident electron energy and *mc*^2^ rest energy of the electron.

For vitreous ice,

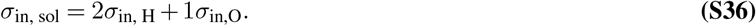

For example, at 300 keV, *σ*_in, sol_ = 343 nm.

#### Phase plate

Based on (36), Eq. S15 can be extended to

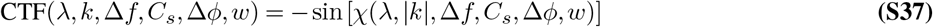

where

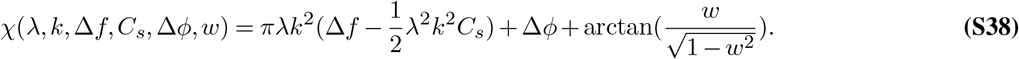

Here, Δ*ϕ* is an additional phase shift introduced by a phase plate and *w* is the fraction of amplitude contrast (e.g., 0.07 or 0.1).

#### Defocus spread from particle thickness

Defocus has variation due to the particle “depth” *D*. We further account for this in the CTF based on (6, 37). For each depth slice *z*, the updated phase shift (Eq. S38) is

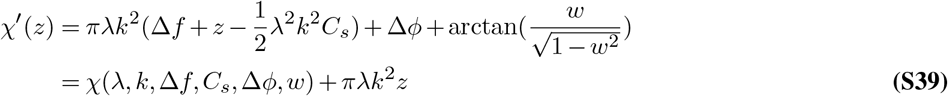

Hence, the depth-averaged CTF is

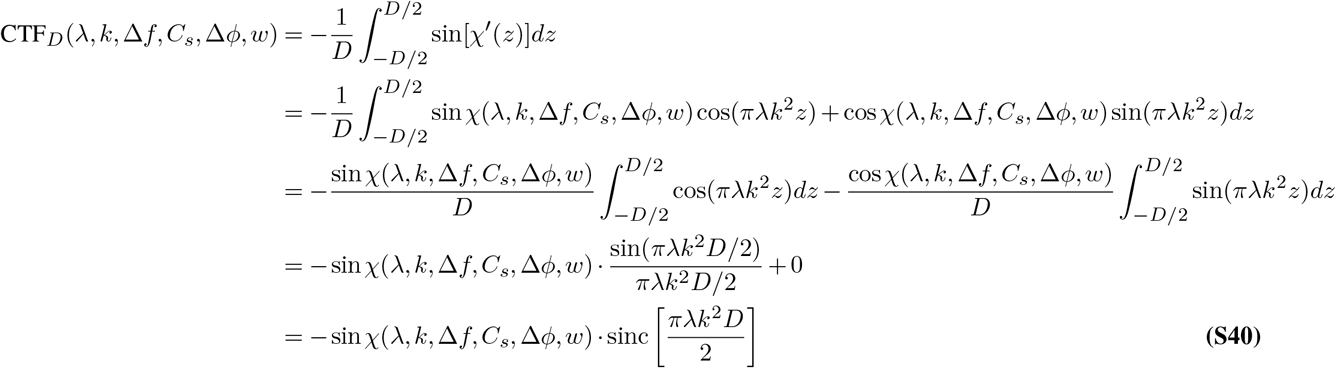

#### Liquid helium cooling

Based on the fact that electron diffraction spots fade 1.2-1.8 *×* slower at liquid helium compared to using liquid nitrogen (24), the exposure filter function can be updated

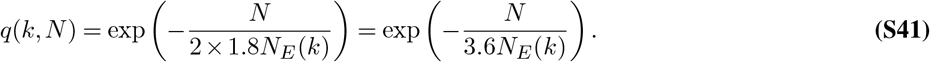

**Figure 1—figure supplement 1.**
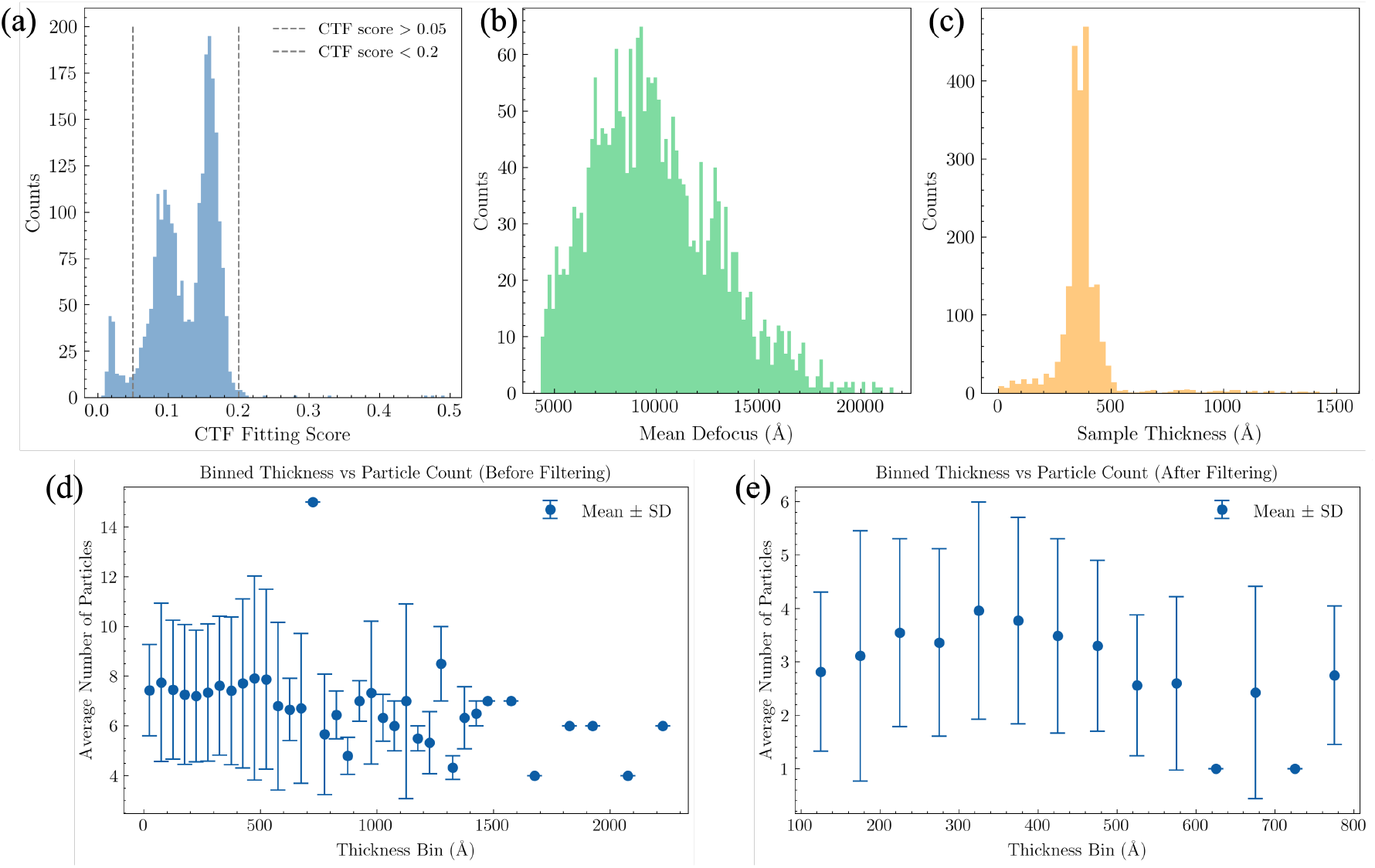
Image statistics of the untilted micrographs in EMPIAR-10252. (a) CTF fitting scores for 2,488 untilted images, calculated using `ctffind5`. Images with scores above 0.2 or below 0.05 were excluded from 2DTM searches. (b) Mean defocus values of the 2,314 images retained for 2DTM. (c) Sample thickness estimates from `ctffind5`, with a median thickness of 367 Å. Negative thickness estimates were excluded from the histogram. (d) Particle counts per thickness bin, based on 17,274 particles extracted from the 2,314 images using `extract-particles`. (e) Particle counts per thickness bin after particle selection using `filter-particles`, showing the final stack of 7,353 particles.

**Figure 1—figure supplement 2.**
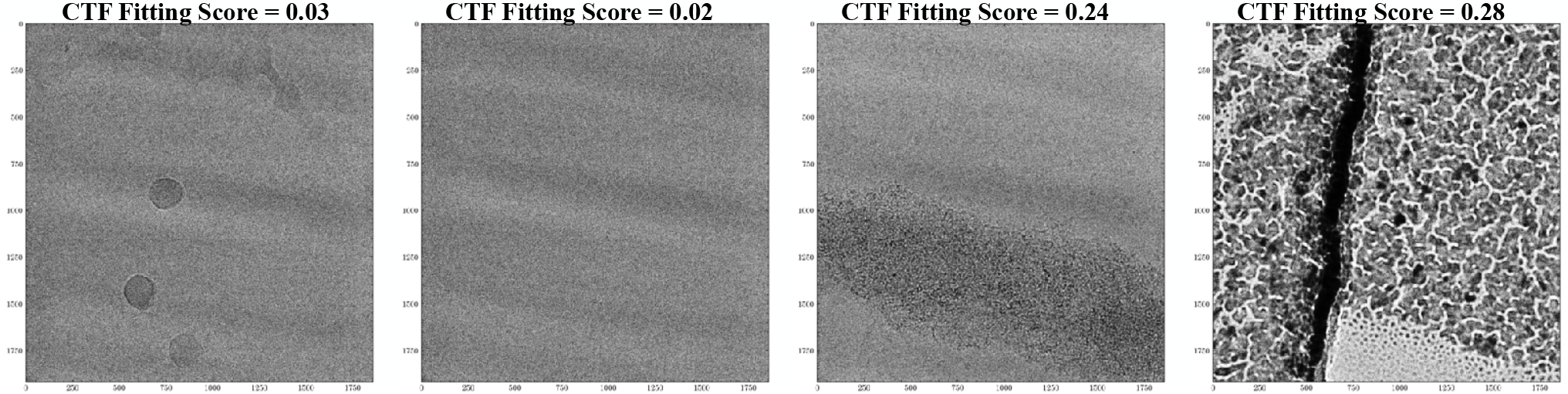
Examples of micrographs excluded from 2DTM search. **(a)** Very low contrast and ice contamination. **(b)** Extremely low contrast, likely drift or astigmatism. **(c)** Particle aggregation or contamination. **(d)** Cross-grating calibration grid.

**Figure 1—figure supplement 3.**
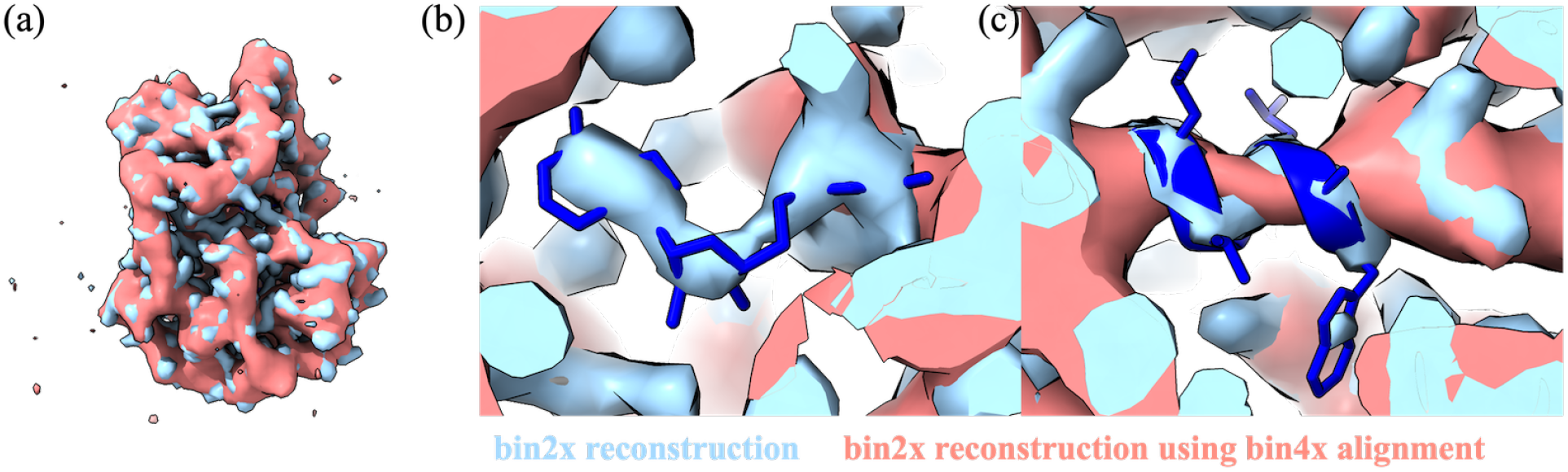
Effect of template resolution on reconstruction quality. Comparison of 3D reconstructions using particles aligned at bin2x (1.117 Å/pixel) versus bin4x (2.234 Å/pixel) resolution. For the bin4x experiment, 2DTM was performed on bin4x images and the detected particle coordinates were used to extract particles from the bin2x images for reconstruction. Densities at the ATP-binding site and deleted residues 222–227 are shown. The bin4x-aligned reconstruction shows loss of ATP density and degraded backbone density for residues 222–227, demonstrating that high-resolution signal in the template is critical for accurate particle alignment and recovery of omitted densities. Map contour *σ* = 5.

**Figure 1—figure supplement 4.**
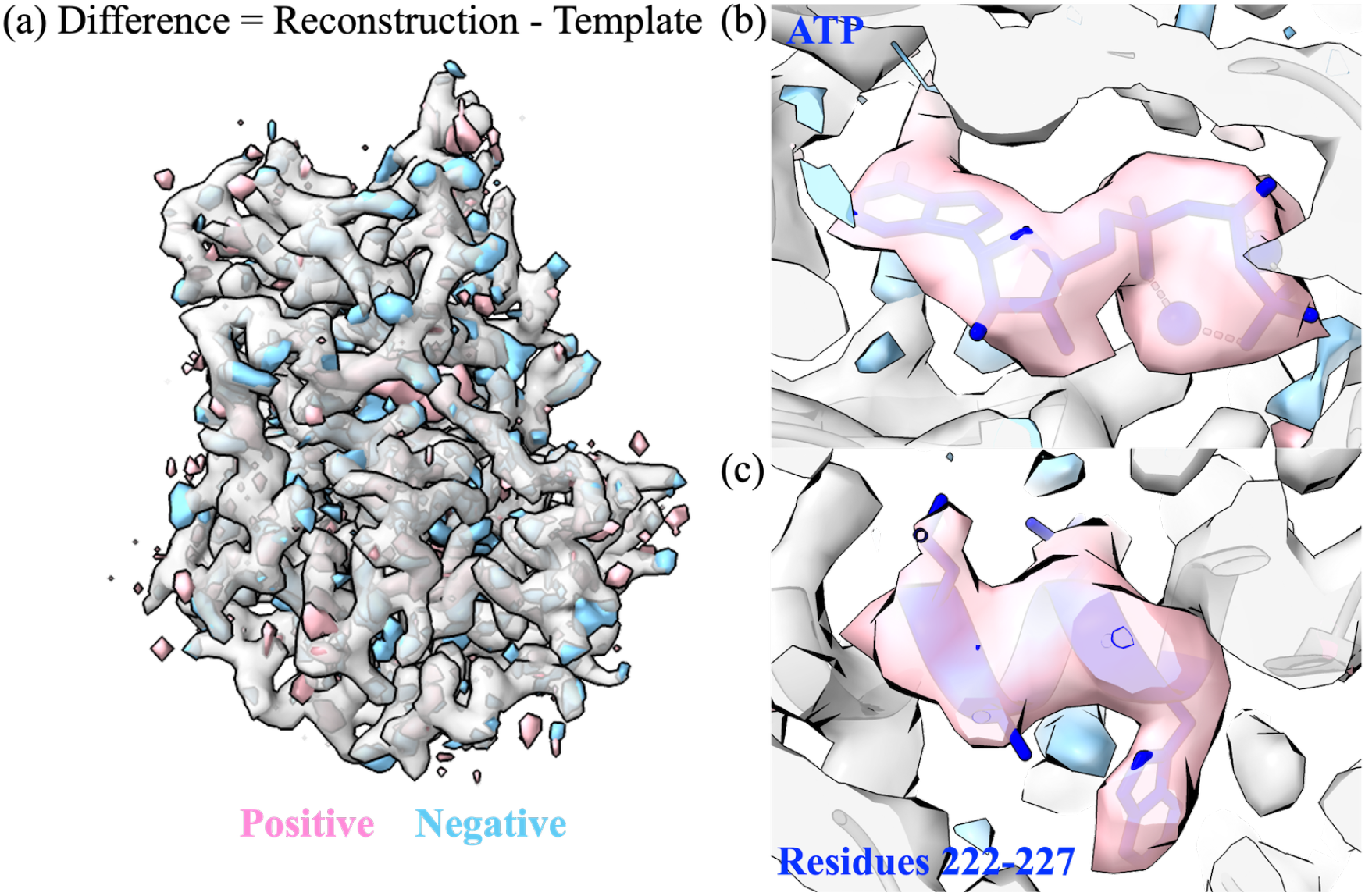
Difference map between the template (gray, map contour *σ* = 15) and the reconstruction shown in Figure 1. The difference map was generated using the `diffmap.exe` program (53) with a protein soft mask. Positive (pink) and negative (light blue) difference densities are shown at map contour *σ* = 20. Positive densities indicate regions present in the reconstruction but absent from the template, with coherent features at the ATP binding site and residues 222–227, which were deleted from the template. Residual noisy densities elsewhere reflect limitations of the forward model.

| Group | Residue | Occupancy |
| --- | --- | --- |
| Omitted residues | Trp222 | 0.70 |
|  | Ala223 | 0.75 |
|  | Leu224 | 0.80 |
|  | Gly225 | 0.55 |
|  | Val226 | 0.70 |
|  | Leu227 | 0.79 |
|  | Mean | 0.72 |
| Ligands | ATP | 0.61 |
|  | Mn<sup>2+</sup> (1) | 0.28 |
|  | Mn<sup>2+</sup> (2) | 0.28 |
| Control residues (in template) | Ile150 | 1.00 |
|  | Val151 | 1.00 |
|  | Leu152 | 1.00 |
|  | Thr153 | 0.89 |
|  | Phe154 | 1.00 |
|  | Glu155 | 0.88 |
|  | Mean | 0.96 |

**Supplementary file 1**. Grouped occupancy refinement of omitted and control residues. Occupancies were refined using Phenix real-space refinement against the omit reconstruction (Figure 1), with the full 1ATP model docked into the map. Omitted residues (222–227), ATP, and Mn^2+^ were absent from the template used for 2DTM alignment. Control residues (150–155) were included in the template. Occupancies near 1.0 for control residues and intermediate values (0.55–0.80) for omitted residues support partial recovery of density in the omitted regions.

|  | p-value = 8.0 | p-value = 7.0 | SNR = 7.5 | + tilt images |
| --- | --- | --- | --- | --- |
| ATP | 0.60 | 0.58 | 0.54 | 0.57 |
| Residues 222–227 | 0.57 | 0.56 | 0.45 | 0.54 |

**Supplementary file 2**. Q-scores for omitted regions across particle selection conditions (Figure 4). Values are average Q-scores over all atoms in each group, calculated using the `MapQ` command line tool (34, 35). The ATP Q-score is averaged over all 31 non-hydrogen atoms; the residues 222–227 Q-score is averaged over all non-hydrogen atoms in the six residues (Trp, Ala, Leu, Gly, Val, Leu).

**Figure 3—figure supplement 1.**
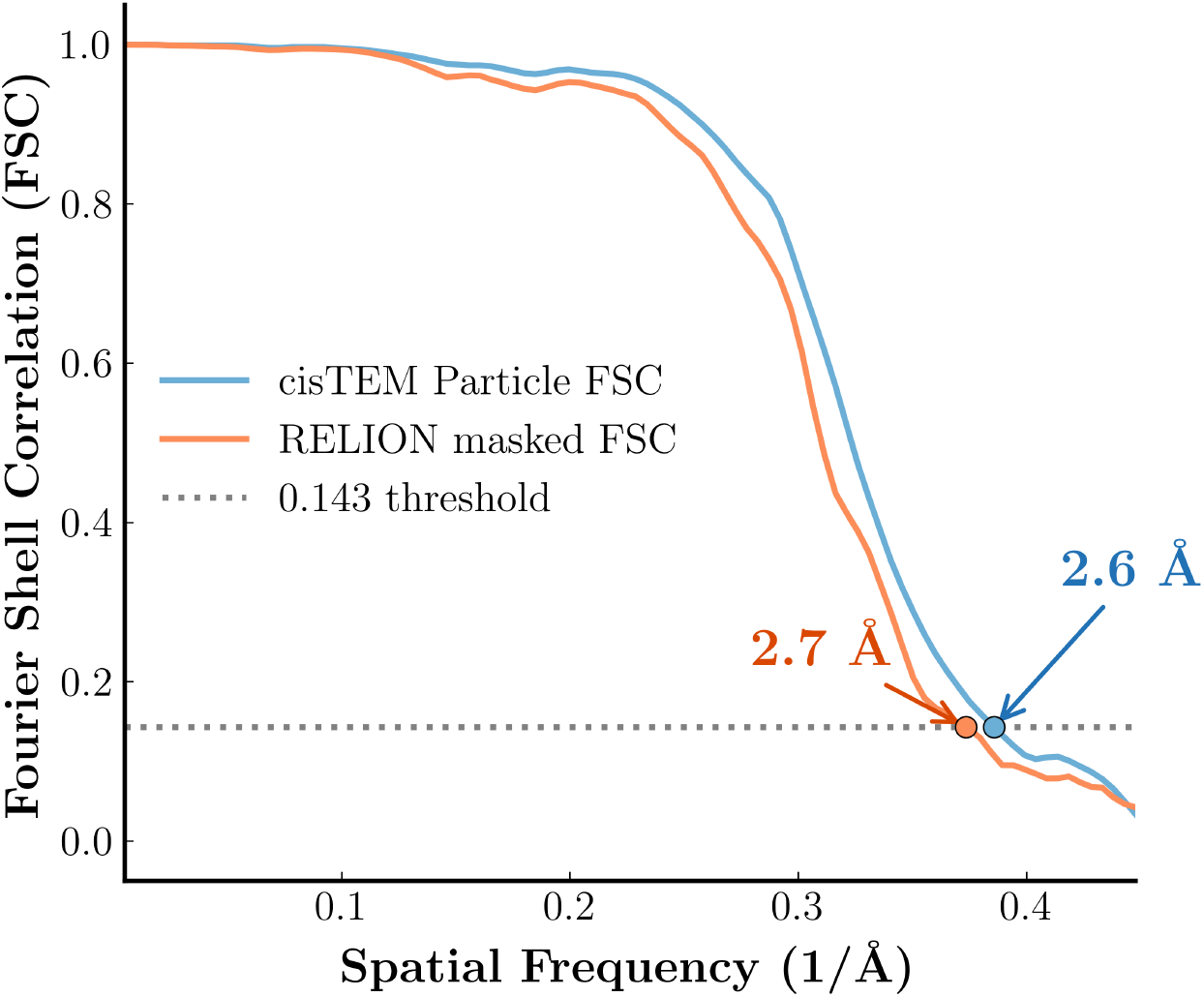
FSC comparison between *cis*TEM and RELION on the same half-maps. Both curves were computed from the same *cis*TEM half-maps (Figure 1 reconstruction). The *cis*TEM Particle FSC (blue) uses a spherical mask with an analytical solvent-fraction correction, while the RELION masked FSC (orange) uses a tight 3D protein mask applied directly to the half-maps. Both cross the FSC = 0.143 threshold at ∼2.6–2.7 Å, confirming that the two packages give consistent resolution estimates when applied to the same data.

**Figure 4—figure supplement 1.**
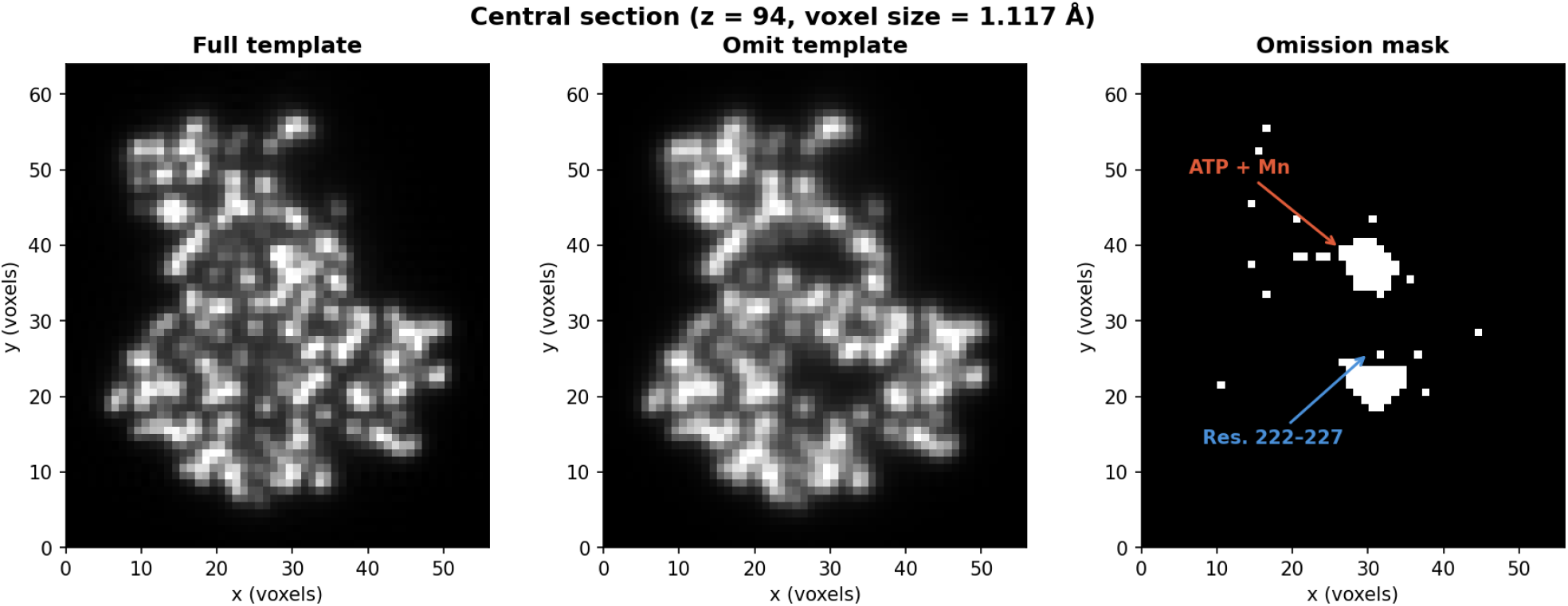
Central section of the omission mask used for template bias (Ω) calculation. The mask was derived from the difference between the full and omit templates (threshold = average of top-100 voxels / 5; 959 masked voxels). Left: full template. Center: omit template (residues 222–227, ATP, Mn, and H2O deleted). Right: binary omission mask, with the two omitted regions (ATP + Mn and residues 222–227) annotated. The mask isolates the omitted region for the Ω calculation shown in Figure 4. Voxel size 1.117 Å.

| Distance from omitted atom (Å) | Omitted | Full occupancy |
| --- | --- | --- |
| 0–1 | +1.84 | +7.77 |
| 1–1.5 | +0.72 | +4.71 |
| 1.5–2 | −0.70 | +0.97 |
| 2–2.5 | −1.41 | −1.23 |
| 2.5–3 | −1.22 | −1.31 |
| *Region average* | −0.38 | +1.65 |

**Supplementary file 3**. Radial profile of the recovered density around omitted residues, averaged across the 36 omit reconstructions. For each reconstruction, voxels in the assigned composite region were binned by distance to the nearest omitted atom. “Omitted” is the mean density in the reconstruction where the residue was absent from the template; “Full occupancy” is the mean of the same voxels in the other 35 reconstructions, in which that residue was present. The atom-centered shells (≤2 Å) recover positive but weakened density upon omission, whereas the peripheral shells (2–3 Å) are negative and nearly identical between the two states. This leads to net-negative density within the molecular envelope (−0.38 versus +1.65 at full occupancy), accounting for the low-resolution negative correlation of the composite map–model FSC (Figure 5—figure supplement 1). Densities are in arbitrary map units.

**Figure 5—figure supplement 1.**
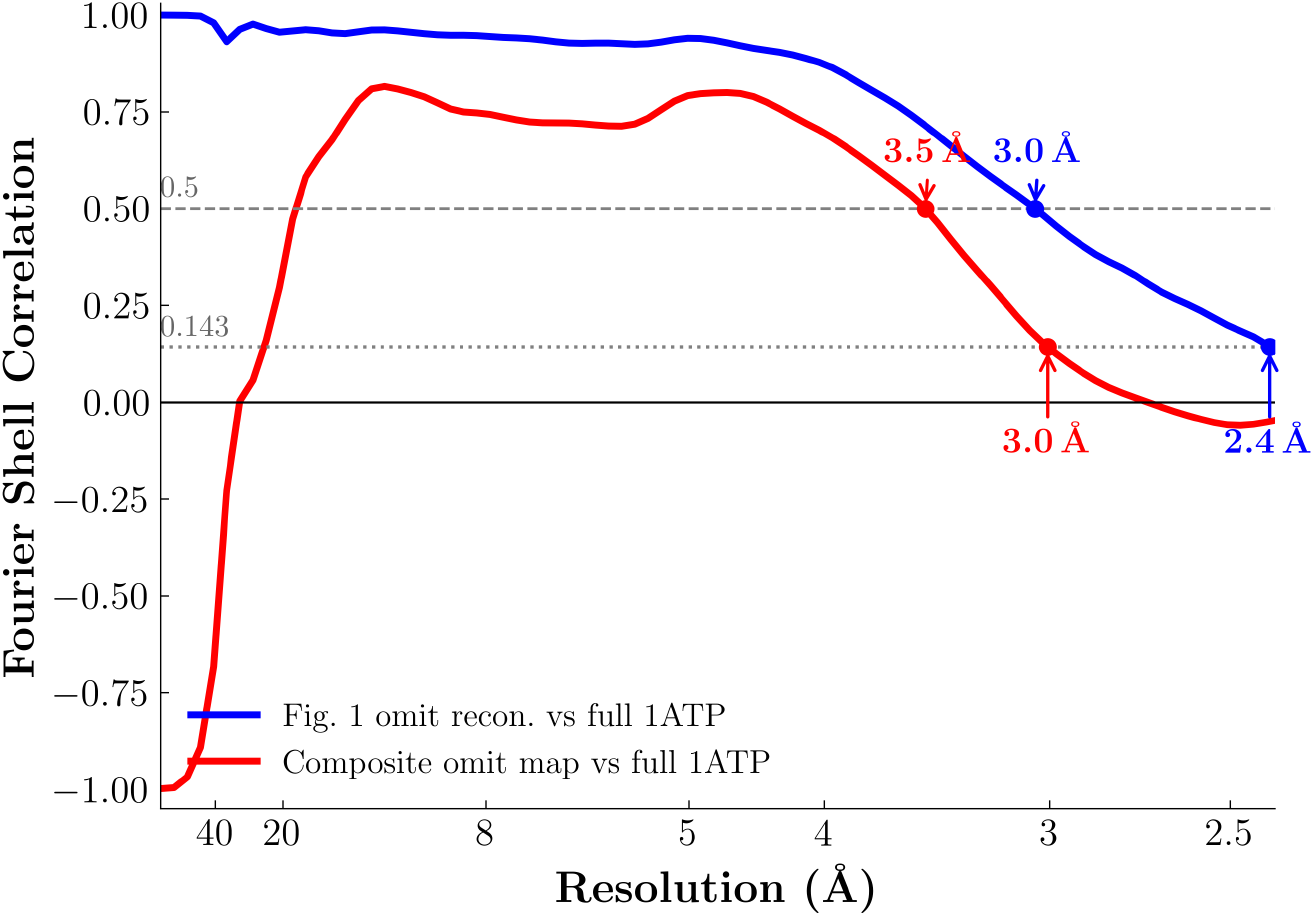
Map–model FSC of the composite omit map (red) and the Figure 1 reconstruction (blue), each computed against a density map simulated from the full 1ATP model using *cis*TEM `calculate_fsc` with a protein envelope mask. Crossings at FSC = 0.5 (dashed) and FSC = 0.143 (dotted) are labelled. The composite omit map crosses FSC = 0.5 at 3.5 Å and FSC = 0.143 at 3.0 Å; the Figure 1 reconstruction crosses at 3.0 Å and 2.4 Å, respectively. Because each voxel of the composite map derives from a reconstruction in which the corresponding residues were absent from the template, the high-resolution agreement reflects genuine signal recovery rather than reproduction of the template. The composite curve falls below the Figure 1 curve because its FSC is dominated by locally omitted density, whereas the Figure 1 curve is dominated by template-included density. The composite curve’s negative correlation at low resolution is examined in Supplementary file 3.

**Figure 5—figure supplement 2.**
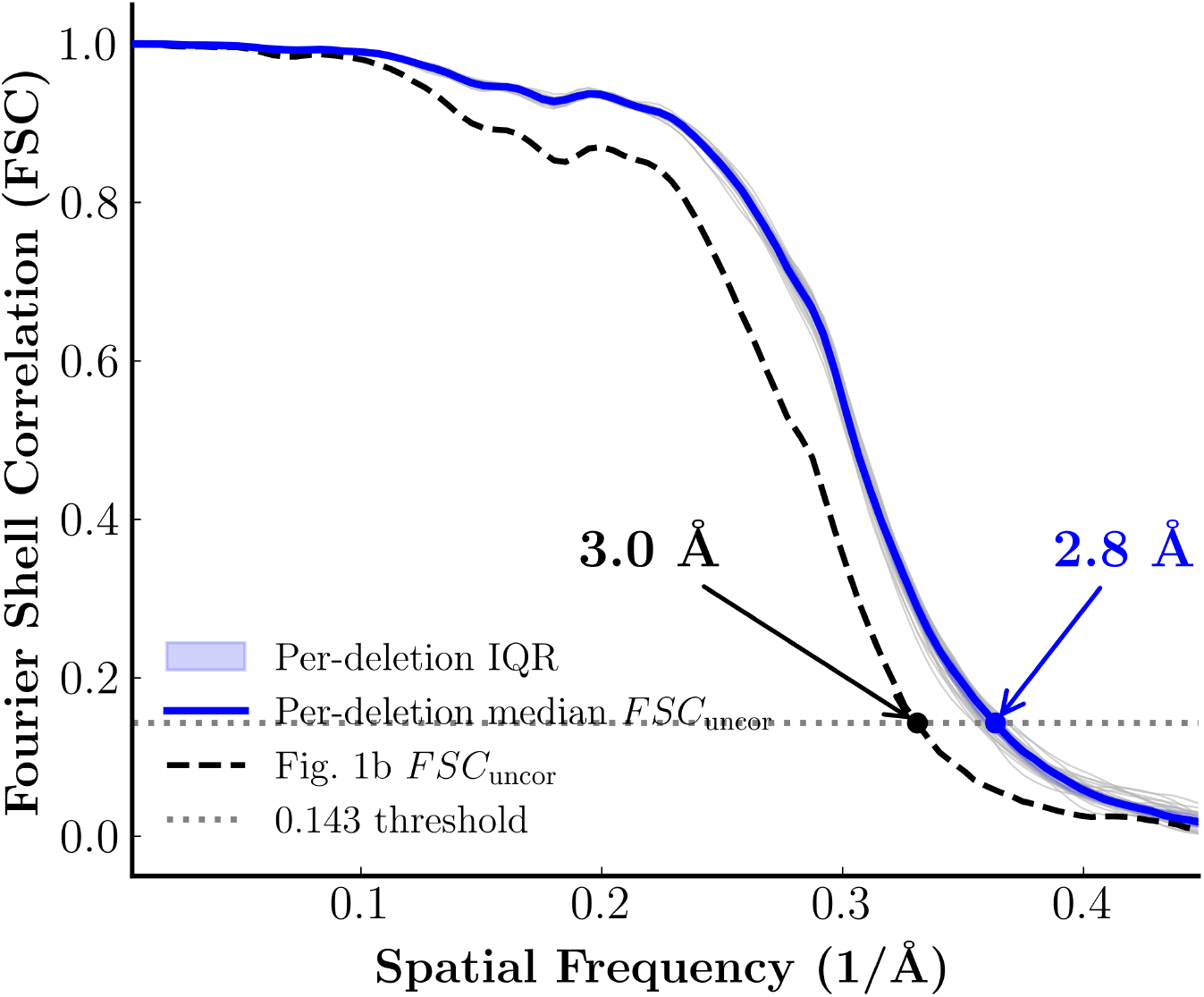
Uncorrected half-map FSC of the 36 individual omit reconstructions (thin grey), with the median (blue) and interquartile range (shaded). The median crosses FSC = 0.143 at 2.8 Å, comparable to the Figure 1b reconstruction (3.0 Å, dashed). Like the global FSC in Figure 1b, these curves use orientations derived from the template and therefore also carry template bias; they are shown for completeness.

**Figure 5—figure supplement 3.**
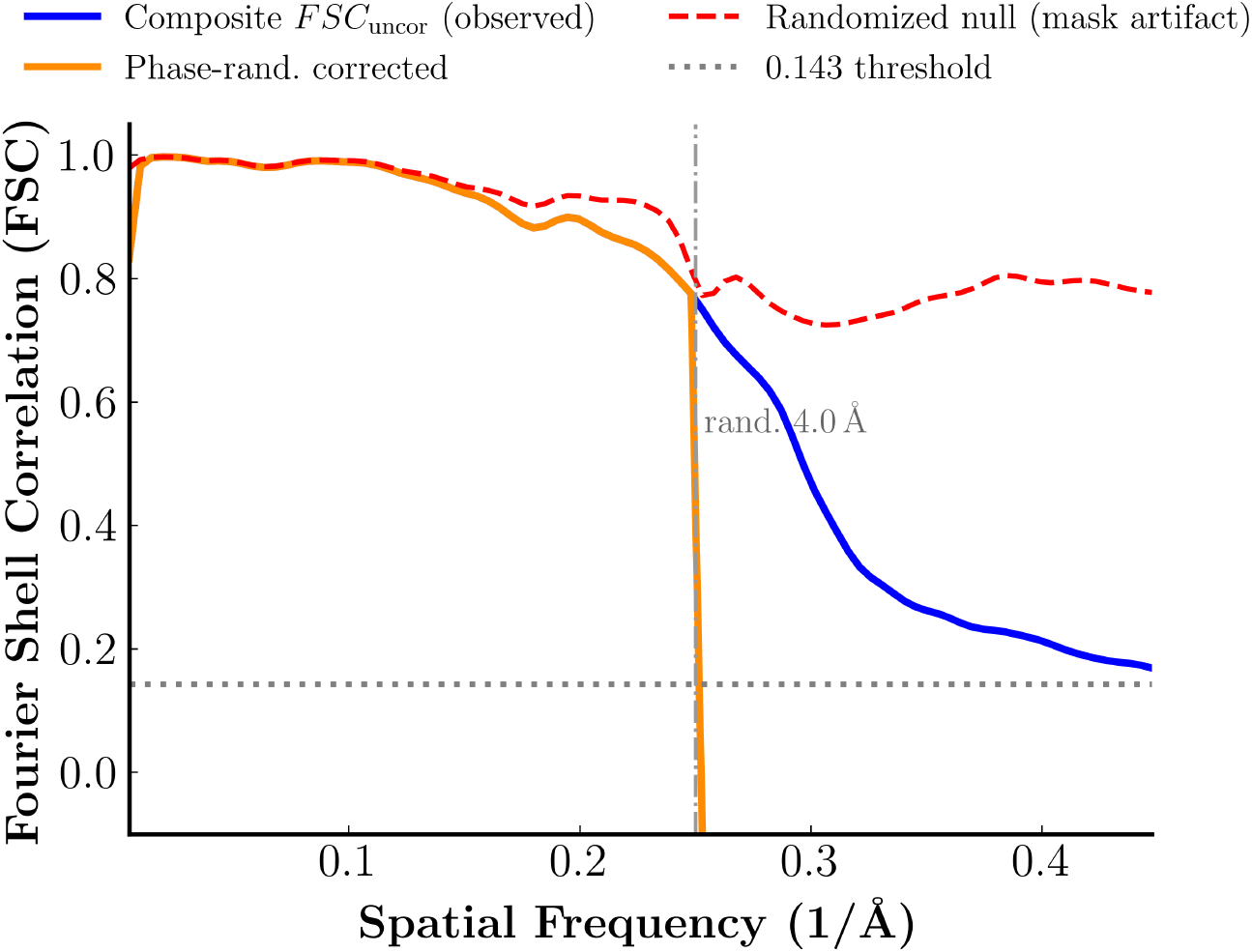
Phase-randomization control for the composite half-map FSC. Shown are the observed FSC between the two composite half-maps (blue), the randomized null (red dashed; the parent half-maps’ phases randomized independently beyond 4 Å and the maps reassembled with the same voxel-assignment masks), and the phase-randomization-corrected FSC (orange). The randomized null remains high beyond the randomization cutoff, indicating that the shared voxel-assignment masks introduce substantial support-induced correlation. Therefore, the observed composite half-map FSC cannot be interpreted as a gold-standard resolution estimate.

**Figure 6—figure supplement 1.**
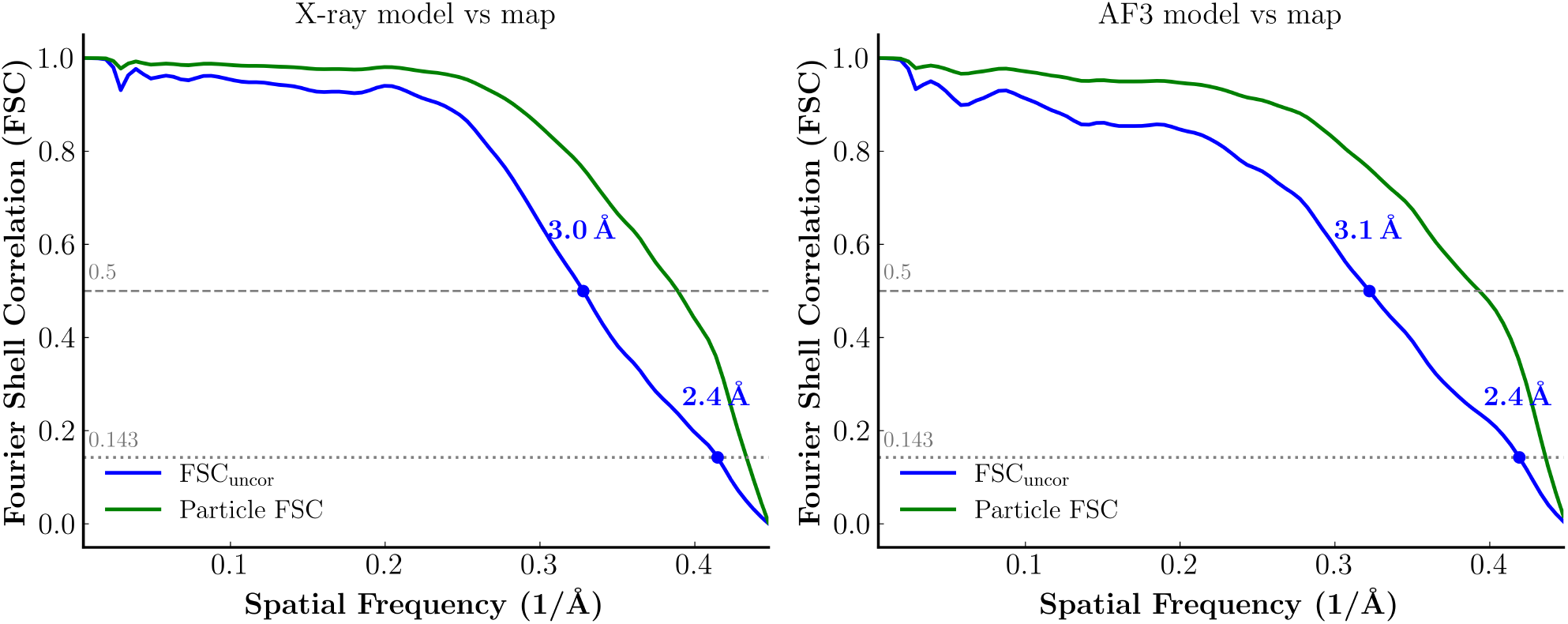
Map–model FSC for X-ray and AlphaFold3 templates in Figure 6. Map–model FSC was computed between each atomic model-derived template and the corresponding 2DTM reconstruction using *cis*TEM `calculate_fsc`, with a protein envelope mask. Left: X-ray-derived template versus X-ray-derived 2DTM reconstruction. Right: AlphaFold3-derived template versus AlphaFold3-derived 2DTM reconstruction. Each panel shows the uncorrected FSC (blue) and the solvent-corrected particle FSC (green); the uncorrected FSC crosses FSC = 0.143 at ∼2.4 Å and FSC = 0.5 at 3.0 Å (X-ray) and 3.1 Å (AlphaFold3).

